# Simulating cell-free chromatin using preclinical models for cancer-specific biomarker discovery

**DOI:** 10.1101/2023.11.16.567416

**Authors:** Steven D. De Michino, Sasha C. Main, Lucas Penny, Robert Kridel, David W. Cescon, Michael M. Hoffman, Mathieu Lupien, Scott V. Bratman

**Author notes:** These authors contributed equally to this work (Steven D. De Michino, Sasha C. Main). **Correspondence to:** Dr. Scott V. Bratman Princess Margaret Cancer Centre, University Health Network, 101 College Street, PMCRT, Room 13-305 Toronto M5G 1L7, Ontario, Canada.

## Abstract

Cell-free chromatin (cf-chromatin) is a rich source of biomarkers across various conditions, including cancer. Tumor-derived circulating cf-chromatin can be profiled for epigenetic features, including nucleosome positioning and histone modifications that govern cell type-specific chromatin conformations. However, the low fractional abundance of tumor-derived cf-chromatin in blood and constrained access to plasma samples pose challenges for epigenetic biomarker discovery. Conditioned media from preclinical tissue culture models could provide an unencumbered source of pure tumor-derived cf-chromatin, but large cf-chromatin complexes from such models do not resemble the nucleosomal structures found predominantly in plasma, thereby limiting the applicability of many analysis techniques. Here, we developed a robust and generalizable framework for simulating cf-chromatin with physiologic nucleosomal distributions using an optimized nuclease treatment. We profiled the resulting nucleosomes by whole genome sequencing and confirmed that inferred nucleosome positioning reflected gene expression and chromatin accessibility patterns specific to the cell type. Compared with plasma, simulated cf-chromatin displayed stronger nucleosome positioning patterns at genomic locations of accessible chromatin from patient tissue. We then utilized simulated cf-chromatin to develop methods for genome-wide profiling of histone post-translational modifications associated with heterochromatin states. Cell-free chromatin immunoprecipitation and sequencing (cf-ChIP-Seq) of H3K27me3 identified heterochromatin domains associated with repressed gene expression, and when combined with H3K4me3 cfChIP-Seq revealed bivalent domains consistent with an intermediate state of transcriptional activity. Combining cfChIP-Seq of both modifications provided more accurate predictions of transcriptional activity from the cell of origin. Altogether, our results demonstrate the broad applicability of preclinical simulated cf-chromatin for epigenetic liquid biopsy biomarker discovery.

## INTRODUCTION

In the past decade, analysis of cancer-specific analytes through a simple blood test, known as liquid biopsy, has emerged as a practice-changing approach for minimally invasive detection of cancer biomarkers^1^. Numerous liquid biopsy analytes have been explored in the context of cancer, most notably circulating tumor cells and cell-free DNA (cfDNA)^1^. cfDNA is released into the bloodstream through various mechanisms — predominantly apoptosis and necrosis — and is digested into small nucleosomal fragments by nucleases^2^. Most cfDNA originates from hematopoietic cells^3^, but in individuals with cancer, a portion of the cfDNA pool consists of tumor-derived fragments that retain the tumor’s genetic and epigenetic alterations. Analysis of tumor-specific cfDNA biomarkers through liquid biopsy provides advantages compared to traditional tissue biopsies, such as overcoming spatial limitations, allowing easily repeatable measurements over time^4^, and increased cost efficiency in certain clinical contexts^5^.

Characterization of tumor-specific genetic alterations within cfDNA has resulted in promising biomarkers for cancer management^6,7^. However, genetic mutations do not consistently reflect the full spectrum of cancer molecular or cellular phenotypes and provide limited information about tumor biology under the influence of anti-cancer therapy^2,8^. Widespread epigenetic aberrations in cancer, such as those impacting DNA methylation, histone modifications, and nucleosome positioning, influence transcriptional activity and cellular phenotypes. The pivotal role of such alterations has motivated the development of new classes of liquid biopsy biomarkers utilizing cell-free chromatin (cf-chromatin)^9^.

Accordingly, emerging methods permit the profiling of epigenetic features in cf-chromatin^2,10–13^. For instance, plasma cf-chromatin is nucleosome bound and protected from nuclease digestion, so it can be leveraged for inferring nucleosome positioning through whole genome sequencing (WGS)^14–16^. This approach resembles the traditional use of nuclease digestion patterns to deduce chromatin structure in cells, such as sequencing protected fragments from micrococcal nuclease (MNase) treatment^17^. Nucleosome positioning inferred from plasma cf-chromatin provides insight into chromatin structure and expression programs from the tissue of origin^14,15,18–20^.

Methods for profiling of histone protein post-translational modifications within cf-chromatin have also recently been introduced. Proof-of-concept studies have demonstrated that histone modifications linked to active (euchromatin)^21^ and repressive (heterochromatin)^22^ chromatin states can be detected within blood plasma. Chromatin states are marked and regulated by a network of histone modifications. Combinations of a core set of these modifications can be used to both discern cell type-specific regulatory programs and predict transcriptional activity^23^. Notably, Sadeh and colleagues profiled cf-chromatin associated with active states through cell-free chromatin immunoprecipitation and sequencing (cfChIP-Seq)^21^; however, sequencing-based profiling of cf-chromatin with histone modifications from repressive chromatin states remains largely unexplored. In cancer, both active and repressive chromatin states are of interest for cell of origin classification, predictive biomarker identification, and resistance mechanism characterization.

A limitation of biomarker discovery for liquid biopsy applications is the heavy reliance on patient plasma samples. For cf-chromatin, this can be particularly problematic due to the <0.1% abundance of tumor-derived material observed in many cancer contexts^24–26^ and the overlapping expression patterns of putative epigenetic biomarkers between tissue types^3,4^. Previous work has attempted to mitigate these issues by selecting samples with high tumor-derived cf-chromatin fractions or by performing deeper sequencing^14,18,19^. While this may be suitable for certain high-burden disease settings, complex biological factors limit the shedding of cf-chromatin from many tumors^27^. Moreover, high cost and low cf-chromatin abundance in finite blood sample volumes present barriers to performing deep sequencing from plasma. Thus, there is demand for models where tumor-derived cf-chromatin can be studied unencumbered by signal dilution and where it is readily accessible and abundant. Preclinical cancer models grown in tissue culture release cf-chromatin that is entirely tumor-derived and easily accessible^28,29^, providing an opportunity to accelerate epigenetic liquid biopsy biomarker discovery and assay development. However, the natural fragment length distribution of conditioned media cf-chromatin contains mainly large fragments (600bp to >10kb), which are less amenable to downstream profiling techniques typically used with plasma, and do not reflect the nucleosomal distributions observed in plasma.

Here, we present a novel framework using a nuclease treatment to simulate cf-chromatin of nucleosomal distributions similar to plasma within conditioned media of preclinical cancer models. This simulated cf-chromatin was amenable to downstream profiling by WGS. We also used simulated cf-chromatin as a renewable source of material to develop for the first time cfChIP-Seq targeting histone modifications associated with both heterochromatin and euchromatin states. We demonstrate that simulated cf-chromatin from media reflects cancer molecular and cellular phenotypes, underscoring its potential utility in liquid biopsy biomarker development.

## METHODS

### Culture models

All cell lines in this study were short tandem repeat (STR) tested to confirm the identity of the parent cell line. All cell lines were grown in their respective media (Supplementary Table S1) supplemented with 10% fetal bovine serum (FBS) and 1% penicillin-streptomycin solution (Wisent Bioproducts). Organoids were cultured in a base medium of Dulbecco’s Modified Eagle Medium/Ham’s F-12 with exogenous growth factors, as previously described^30^. All cell lines were mycoplasma tested prior to freezing down stocks and prior to experiments using the e-Myco™ VALiD Mycoplasma PCR Detection Kit (iNtRON Biotechnology, Inc., CAT #25239).

### Media sample collection and processing

Media samples were collected in the exponential growth phase, approximately three to four days after seeding. Media was placed in 15 mL conical tubes and spun at 2500g for 10 min at 4 °C. Media was transferred to centrifuge-safe 5 mL Eppendorf tubes and spun at 16100g for 10 min at 4 °C. The supernatant was transferred to a new 15 mL conical tube and kept at 4 °C until nucleosome preparation without freezing.

### Quantification of cf-chromatin in culture media

To determine the concentration of cf-chromatin in the media of each cell line, DNA was quantified using quantitative polymerase chain reaction (qPCR) against human LINE-1^31,32^ in quadruplicate. Media was diluted 1:100 in low TE DNA suspension buffer (10 mM Tris-HCl, 0.1 mM EDTA, pH 8.0, TEKnova, CAT #T0220). Human male genomic DNA (Promega, CAT #G1471) was used for a standard curve at 125 pg/µL, 12.5 pg/µL, 1.25 pg/µL, 0.125 pg/µL, and 0.0125 pg/µL. Each 10 µL reaction consisted of 4 µL template, 2x SsoAdvanced Universal SYBR® Green Supermix (BioRad, CAT #1725274), and 2.5 µM oligonucleotides targeting hLINE-1 (Supplementary Table S2).

### Nucleosome preparation with MNase

For nucleosome preparation from cell culture media, 1–5 mL conditioned media was subjected to digestion with a fixed concentration of MNase. MNase master mix was prepared at 3000 Units/mL (U/mL) by combining 440 μL of ddH20, 50 μL of MNase buffer, 5 μL of BSA, and 5 μL of 300,000 U/mL MNase (New England Biolabs, CAT #M0247S). Efforts were made to avoid freeze-thaw cycles of the MNase enzyme. MNase was added to cell line conditioned media supernatant at a final concentration of 20 U/mL and incubated in a 37 °C water bath for 30 min. The reaction was stopped by adding 5 mM EDTA (Invitrogen, CAT #15575020).

### cfMNase-Seq

Following nucleosome preparation of media cf-chromatin, purification of DNA from cf-chromatin was performed using the Qiagen Circulating Nucleic Acid kit (CAT #55114). Purified nucleosomal DNA from cf-chromatin was quantified using the Qubit High Sensitivity dsDNA kit (Life Technologies, CAT #Q33231) according to kit instructions. DNA fragment length distributions were analyzed on an Agilent 2100 Bioanalyzer with the High Sensitivity DNA kit (CAT #5067-4626) according to the manufacturer’s instructions. Purified DNA was stored at either 4°C (<3 days) or −20°C (>3 days). Illumina-compatible sequencing libraries were prepared from 20 ng of purified DNA in 50 μL of TE buffer using the KAPA Hyperprep Kit with Library Amplification Primer Mix (Roche, CAT #07962363001) and 6 cycles of PCR amplification. Sequencing adapters containing unique molecular identifiers (UMIs) were obtained from Integrated DNA Technologies^33^. Library clean-ups were performed using Beckman Coulter AMPure XP beads (CAT #A63882) with a 0.8× cleanup before library amplification and a 0.9X cleanup afterward. The library concentration was determined using the Qubit High Sensitivity dsDNA kit and the fragment length distribution was assessed with an Agilent 2100 Bioanalyzer. All cfMNase-Seq samples were then pooled and sequenced by Illumina NovaSeq (41.2-69.2 million (M), 100 bp paired-end (PE) reads, mean 52.4M).

### cfChIP-Seq

Following nucleosome preparation of media cf-chromatin, a predetermined quantity (10-300 ng SU-DHL-6 cfDNA) was diluted in RPMI 1640 (Wisent Bioproducts, CAT #350-000-CL) and subjected to ChIP-Seq adapted from Sadeh et al^21^. Monoclonal antibodies targeting H3K4me3 (Abcam, ab1012) and H3K27me3 (Cell Signaling Technology, 9733S, lot 21) were first validated for antigen specificity by Western blot using recombinant H3K4me1, H3K4me2, H3K4me3, and H3K27me3. For the Western blots, 0.5 µg of recombinant histone was loaded per lane, and samples were run on a 4-20% polyacrylamide gradient gel (Bio-Rad, CAT #4568094) for 40 min at 100 V. Semi-dry transfer was utilized, and 1:1000 and 1:10000 dilutions of primary and secondary (Li-cor, CAT #925-32211/925-32210) antibodies, respectively, were used for detection.

To block any non-specific chromatin absorption during ChIP, Dynabeads (Invitrogen, CAT #14301) conjugated with antibodies were pre-incubated with 5% bovine serum albumin (BSA) in ddH2O for 1 h at room temperature. After one wash with media (without FBS) to remove residual BSA, the media sample and 3 μg dynabead-conjugated antibody were incubated overnight at 4°C. Beads were then washed prior to conducting on-bead library preparation as described^21^ using the KAPA Hyperprep Kit with Library Amplification Primer Mix. H3K4me3 libraries were amplified to 16 cycles, and H3K27me3 libraries were amplified to 12 cycles for 10-30 ng input and 11 cycles for 300 ng input. Library clean-ups were performed using AMPure XP beads with a 0.8X cleanup before library amplification and a 0.9X cleanup afterward. Library concentration was measured with the Qubit High Sensitivity dsDNA kit; fragment size distribution was evaluated by the Agilent 2100 Bioanalyzer. Samples were pooled and sequenced by Illumina NovaSeq (8.04-12.4M, mean 9.79M, 100PE for H3K4me3 libraries; 107.8-163.6M, mean 129.6M, 100PE for H3K27me3 libraries).

### ATAC-Seq

Assay for transposase-accessible chromatin using sequencing (ATAC-Seq) data for the cell line CAMA-1 was generated following a similar protocol as previously described^34^. Briefly, two biological replicates of 50,000 cells were lysed and used for the transposition reaction. An in-house Tn5 enzyme was used for the transposition reaction, prepared as previously described^35^. Sample clean-up was performed with the Qiagen MinElute PCR Purification Kit (CAT #28004) with 250 μL of buffer PB for each 50 μL sample. A qPCR was then performed to determine the number of PCR amplification cycles to perform, and then the libraries were amplified (7-8 cycles). The samples were then cleaned up with the Qiagen MinElute PCR Purification Kit, diluted in 30 μL, and size selected with AMpure XP beads (0.5X). Lastly, a quality control qPCR was performed using 2.5 μL of diluted sample (6 μL sample in 30 μL water) and 7.5 μL of master mix (5 μL of 2X SYBR, 1 μL of 5 µM forward and reverse primers, and 2 µL of water) for two constitutively open (*GAPDH* and *KAT6B*) and closed regions (*QML93* and *SLC22A3*) (Supplementary Table S2). All samples had fold enrichment greater than ten and were considered acceptable. The libraries were then quantified using a Qubit High Sensitivity dsDNA kit, and fragment length distribution was assessed with an Agilent 2100 Bioanalyzer. Samples were then pooled and sequenced by Illumina NovaSeq (90.4-103.9M, mean 97.2M, 100PE).

### RNA-Seq

RNA was extracted from CAMA-1 cells with NucleoSpin RNA kit (Machery Nagel). PolyA sequencing libraries were prepared and RNA-Seq was conducted by Novogene Corporation (40M reads, Illumina 150PE).

### Sequence data processing

#### Cf-chromatin data

Paired-end sequencing reads were trimmed using Cutadapt (v3.0)^36^, utilizing a list of UMIs as input for cfMNase-Seq and cfChIP-Seq samples. Trimmed reads were aligned to the hg38 human genome build using Bowtie2 (v2.4.1)^37^. Duplicate reads were removed using Samtools (v1.9)^38^, using a combination of samtools sort (before and after samtools fixmate), samtools fixmate with the -m option, and samtools markdup with the -r option. Low-quality mapped reads were removed using samtools view with the -bq option, with a minimum mapping quality of 25-30. Fragment length distribution was assessed by Picard CollectInsertSizeMetrics (v2.6.0), and coverage was assessed with plotCoverage from DeepTools (v3.2.1)^39^. Counts per 50 bp bin were calculated from the output BAM files (in bigWig format), using DeepTools^39^. For cfMNase-Seq samples, bigWig files were generated with Reads Per Kilobase per Million mapped reads (RPKM) normalization, centered, and extended read options. For cfChIP-Seq samples, profiles that were not subsampled down for peak comparison (for instance, H3K4me3 in Figure 3D) were RPKM normalized in bigWig format. Peaks were called using MACS2 (v2.2.7.1)^40^ using the narrow peak option for H3K4me3 and the broad peak option for H3K27me3. ENCODE blacklist regions^41^ were removed from peak sets and counts files.

#### ATAC-Seq

Reads were processed in a similar method as above using a predefined pipeline (https://github.com/LupienLab/pipeline-chromatin-accessibility). Briefly, TrimGalore (v0.6.5) was used to trim FASTQ files, Bowtie2 (v2.3.5.1) was used for alignment, and MACS2 (v2.2.6) was used for peak calling. The analytical pipeline is also made reusable on the CoBE platform (www.pmcobe.ca/pipeline/60a4336aaf7a3251ac7e152d).

#### RNA-Seq

Expression matrices were constructed via a custom pipeline (https://github.com/elsamah/rna-Seq-star-deseq2). Briefly, FASTQ files were aligned to hg38 using STARv2 (v2.7.8a)^42^ aligner, and gene expression levels were quantified using RSEM (v1.3.0)^43^. Fragments per kilobase of transcript per million mapped reads (FPKM) and transcripts per million (TPM) normalized counts were used for downstream analyses. SU-DHL-6 RNA-Seq^44^ was processed using TrimGalore (v0.6.6) for trimming (https://www.bioinformatics.babraham.ac.uk/projects/trim_galore/https://github.com/FelixKrueger/TrimGalore, implementable wrapper for CutAdapt^36^), TopHat2^45^ for alignment to GRCh38/hg38^46^. Next, Samtools^38^ for removing duplicate reads, and deepTools to obtain RPKM normalized counts per 50 bp bin.

### Genome-wide cfMNase-Seq and MNase-Seq analysis

BigWig files with a 10,000 bp bin size were generated for all cfMNase-Seq samples in deepTools^39^, uploaded to the CyVerse Discovery Environment^47^, and visualized in the UCSC Genome Browser^48^ as a custom track. Next, the average scores for each bigWig file were computed across every genomic region using the multiBigWigSummary command in deepTools^39^, producing a compressed NumPy array. A Spearman correlation was then performed on the array for all cfMNase-Seq samples and visualized with the command plotCorrelation in deepTools^39^. Next, a principal component analysis (PCA) was performed on all cfMNase-Seq and MNase-Seq samples using the plotPCA command in deepTools^39^, and was replotted in R (v4.2.1) using ggplot2 (v3.4.0)^49^. K-means clustering was performed using R-base packages, and a silhouette score was calculated per clustered group using Cluster (v2.1.4). The silhouette score is calculated as follows: for any point *i:*

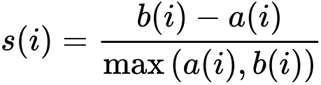

Where *a(i)* represents the average intra-cluster distance between *i* and all other points in the cluster and *b(i)* represents the inter-cluster distance between *i* and the nearest cluster centroid; the average silhouette score is calculated for all data points within the dataset.

### cfMNase-Seq copy number aberration analysis

Copy number aberration analysis was conducted with HMMcopy readCounter (https://github.com/shahcompbio/hmmcopy_utils)^50^ and ichorCNA (v0.2.0)^51^. The analysis was run using the default parameters and default panel of healthy normal samples, except that we limited the analysis to autosomes and decreased the set of non-tumor fraction parameter restart values to (0.001,0.01,0.1,0.15,0.2,0.25) since our cf-chromatin samples are entirely tumor-derived DNA. The default optimal solution was reported from this analysis.

### Reference file generation and cfMNase-Seq coverage analysis

#### Gene expression reference files

Gene expression reference files were created using RNA-Seq data for each respective cell line by subsetting transcripts by FPKM level (FPKM=0, 0<FPKM≤0.1, 0.1<FPKM≤1, 1<FPKM≤10, FPKM>10).

#### Chromatin variant reference files

The ATAC-Seq BAM and narrowPeak files from HCT116 and CAMA-1 (two replicates each) were used as input into diffBind (v3.6.5)^52^ to identify sites of differential chromatin accessibility. Analysis followed methods previously described^52,53^, and all parameters were set to defaults, except for normalization, which was set to background (https://bioconductor.org/packages/devel/bioc/vignettes/DiffBind/inst/doc/DiffBind.pdf).

DESeq2 (v1.36.0)^54^ within diffBind was used to select differential sites, and those with a false discovery rate less than 0.01 and a log2 fold change greater than two or less than negative two were considered differential. The distribution across genomic elements of each reference file was plotted using peak annotation within ChIPseeker (v1.32.1)^55^. Reference site files were saved separately for differential open chromatin sites enriched in CAMA-1 (81,292 regions) and HCT116 (19,807 regions).

#### Cancer patient chromatin accessibility reference files

Open chromatin sites for primary breast cancer and colon adenocarcinoma patient samples from The Cancer Genome Atlas (TCGA) were downloaded from the TCGA ATAC-Seq hub (https://gdc.cancer.gov/about-data/publications/ATACseq-AWG)^56^. A file with all cancer type-specific peak sets was downloaded (https://api.gdc.cancer.gov/data/71ccfc55-b428-4a04-bb5a-227f7f3bf91c), and files for breast cancer (BRCA_peakCalls.txt) and colon cancer (COAD_peakCalls.txt) were obtained.

#### Coverage analysis

All reference files were formatted to be used as input site files for coverage evaluation with Griffin (v0.1.0)^15^. Griffin was run for cfMNase-Seq (n=6 for CAMA-1, n=6 for HCT116) and patient plasma WGS samples (n=27 for colorectal cancer, n=54 for breast cancer)^57^ with all default parameters, except for the individual option, which was set as true. The coverage profiles and quantitative metrics were then plotted in R with ggplot2^49^.

### cfMNase-Seq and ATAC-Seq comparison

A reference track file was downloaded from UCSC Table Browser^58^ with TSSs for all known genes. BigWig files were generated for ATAC-Seq data and scores across TSSs were calculated using deepTools^39^. The Griffin^15^ pipeline was then modified to extract G+C-corrected bigWig files for cfMNase-Seq coverage across the entire length of all fragments (as opposed to midpoint coverage for fragments 100-200 bp). The ATAC-Seq signal sorted region files were then used to plot heatmaps in deepTools.

### cfChIP-Seq analysis

SU-DHL-6 media cf-chromatin H3K4me3 profiles were visualized over TSSs using deepTools^39^ (v3.2.1), with the computeMatrix (reference-point, -a 3000, -b 3000 – skipZeros, and –missingDataAsZero options) and plotHeatmap functions. Correlation matrices were generated in deepTools using the multiBigwigSummary and plotCorrelation functions with either -pearson or -spearman options.

All analyses listed below using R statistical software were performed using R (v4.1.1). Data transformation was done using the Tidyverse suite (v1.3.1), including ggplot2 (v3.4.1) for plotting. Stacked BioAnalyzer traces were re-plotted using ggplot2, using the raw output from the Agilent BioAnalyzer 2100 (v.B.02.10.SI764). All statistics including t-tests were performed using the R stats package. Peak overlap analysis and Venn diagrams were done using ChIPPeakAnno (v3.28.0). UpSet plot was generated using UpSetR (v1.4.0), using the output from HOMER mergePeaks (https://github.com/stevekm/Bioinformatics/blob/776c420efac851c6780ce573939fb6610a3b9ae8/HOMER_mergePeaks_pipeline/HOMER_mergePeaks_venn_UpSetR/multi_peaks_UpSet_plot.R). HOMER mergePeaks was used to find the overlap of multiple H3K4me3 peak sets, displayed using the UpSet plot.

Unsupervised hierarchical clustering was performed using the stats package (R v4.1.1)^59^ and dendextend (v1.16.0)^60^. bedtoolsr (v2.30.0-4)^61^ was used to sort features from the custom heatmap in the H3K4me3 peak analysis. BiomaRt (v2.50.0)^62^ was used to generate the TSS reference for the custom heatmap. Color palettes were primarily derived from RColorBrewer (v1.1-3)^63^ and MetBrewer (v0.2.0)^64^. ChIP-Enrich (v2.18.0)^65^ was used to find the association between H3K4me3 peaks and active cellular pathways.

Differential histone modification analysis was performed using DiffBind (v3.8.4)^53^ in R (v4.2.1) using standard parameters. SU-DHL-6 media and OCILY3 nuclear ChIP-Seq samples were grouped into biological replicates and quality assessment of ChIP-Seq data was performed using ChIPQC (v1.34.1)^66^. ChIP-Seq peaksets were read using the DiffBind function dba. Next, reads were counted and normalized using dba.count and dba.normalize, respectively. Differential peaks were computed using dba.analyze against OCILY3 samples with 2 minimum members and differentially bound sites were retrieved using dba.report. Finally, high-confidence differentially bound peaks were annotated and visualized using ChIPseeker (v1.35.3)^55^ and TxDb.Hsapiens.UCSC.hg38.knownGene (v3.15.0)^67^.

### Differential mRNA Expression Analysis

Differential mRNA Expression Analysis was performed with DESeq2 (v1.36.0)^54^ using standard parameters in R (v4.2.1). Gene expression counts were computed as described earlier with GRCh38/hg38^46^. Differential scores were calculated for GCB DLBCL against ACB DLBCL subtype and filtered for a minimum expression level of log_2_ ≥ 1.2 and FDR <0.05. This resulted in 314 upregulated and 1085 downregulated genes.

### Supervised Binary Classification of SU-DHL-6 Histone Modifications

Binary classification of SU-DHL-6 gene expression from histone mark modifications was performed using an adapted version of ShallowChrome from Frasca et al.^68^ to predict gene expression. 56 cell types from Roadmap Epigenomics Consortium^23^, 9 cell types from ENCODE^69–72^, and a lab-generated cfChIP-Seq SU-DHL-6 media sample, each of which had paired histone modification and gene expression data, were used for this analysis and all were processed under ChIP-Seq uniform processing guidelines as previously described^23^ (Supplementary Table S3). This approach used peak calling within each gene’s TSS and gene body to predict gene expression for each given cell type. We applied 10-fold cross-validation of each cell type where approximately 6000 genes were held out in each fold and performance was computed using AUC for each individual gene feature and region combination.

## RESULTS

### Nuclease treatment of cell line conditioned media reproducibly generates nucleosomal distributions from cf-chromatin

To address the limitations of current cf-chromatin liquid biopsy discovery tools, we devised a framework for simulating cell-free nucleosomes using tissue culture conditioned media (Figure 1A). First, we collected media from cultured HCT116 (colon cancer) cells during log-phase growth and measured cf-chromatin topology by DNA fragment size analysis (Figure 1B). In stark contrast to the mono- and oligo-nucleosomal topology of cf-chromatin present in human plasma^14^, this native untreated cf-chromatin contained predominantly DNA fragments larger than 600 bp in length (median ∼3,000 bp).

**Figure 1.**
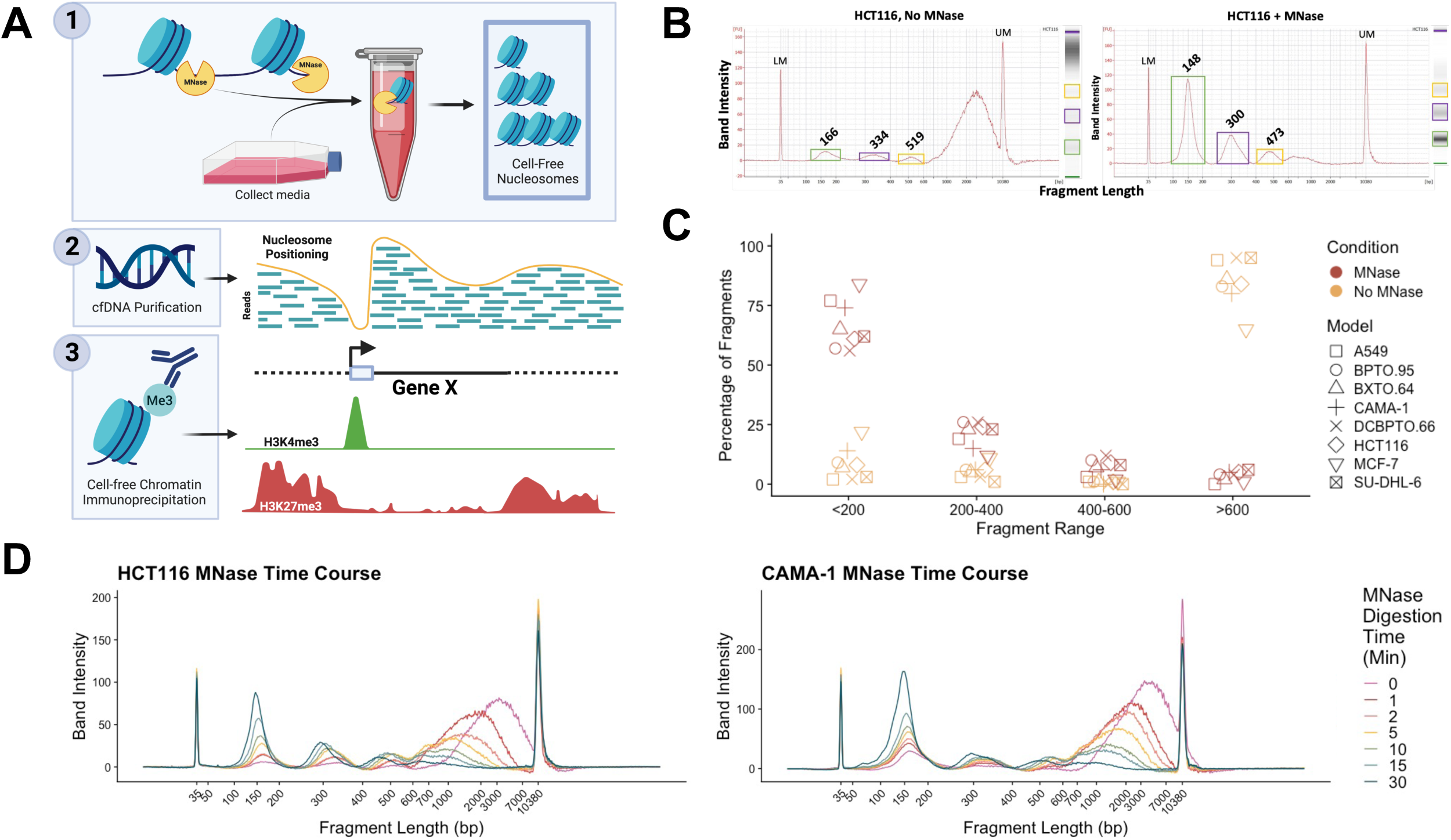
MNase treatment of media from preclinical models reproducibly generates nucleosomal distributions from cf-chromatin. **A)** Graphical abstract of simulated cf-chromatin and downstream profiling techniques. **B)** BioAnalyzer traces showing DNA fragment length distribution from simulated cf-chromatin. Compared with untreated HCT116 conditioned media (left), MNase treatment (right) leads to enrichment for mono- and oligo-nucleosome-sized cfDNA fragments. LM=lower marker, UM=upper marker. **C)** MNase treatment generates significantly different proportions of mono-di- and tri-cf-nucleosomes compared to no nuclease treatment, reproducible across various 2-D and organoid culture models (CAMA-1 and MCF7=breast adenocarcinoma, HCT116=colorectal carcinoma, A549=lung adenocarcinoma, SU-DHL-6=diffuse large B-cell lymphoma, BPTO.95 and DCBPTO.66=breast cancer patient tumor-derived organoids, BXTO.64=breast cancer patient-derived xenograft derived organoid). Paired t-test for fragments: <200 bp, p=1.53×10^-7^; 200-400 bp, p=5.05×10^-4^; 400-600 bp, p=1.44×10^-3^; >600 bp, p=6.07×10^-8^. Data are derived from the BioAnalyzer traces shown in Supplementary Figure S1B. **D)** MNase treatment time course of HCT116 and CAMA-1 conditioned media. Samples were treated with MNase over a 0 to 30-minute time course. BioAnalyzer traces show an increasing proportion of mononucleosome-sized fragments with longer digestion time.

To generate cf-chromatin with more physiological properties reflective of human plasma, we subjected HCT116 conditioned media to nuclease digestion with MNase. MNase is commonly used in “native” ChIP-Seq experiments in place of sonication to avoid disturbing transcription factor association with DNA^73^, in Cleavage Under Targets and Release Using Nuclease (CUT&RUN)^74^, and for mapping nucleosome occupancy^2,14^. Following 30 minutes of digestion, we observed a striking shift from large fragments to smaller fragments corresponding to the size of mono-, di-, and tri-nucleosomes, with the median size of the mononucleosomal DNA fragments being 148 bp. This characteristic size distribution was not fully maintained in culture media that was subjected to freeze-thaw (Supplementary Fig. S1A), indicating that fresh media is preferred for optimal nucleosome integrity.

### Simulated cf-chromatin is robustly produced across tissue culture models

Having demonstrated the ability of MNase treatment to produce mono- and oligo-nucleosomal populations of cf-chromatin within HCT116 cultured media, we next sought to evaluate the generalizability of this technique in other preclinical models. We observed a significant difference in the fragment length distributions between conditions across various 2D and 3D tissue culture models (Figure 1C, Supplementary Fig. S1B), with the nuclease treatment condition better capturing nucleosomal size distributions observed in human plasma than media cf-chromatin without nuclease treatment. Across all tested models, significant enrichment in fragments greater than 600 bp was observed in untreated culture conditions, while fragments less than 200 bp were most enriched in the MNase treatment condition. Recognizing the intrinsic variability in the amount of cf-chromatin released among distinct models^28,29^, we confirmed consistent nucleosome size distributions across a range of cf-chromatin input quantities and volumes (Supplementary Fig. S1C).

Maintenance of the characteristic mononucleosomal size distribution within cf-chromatin was strongly time-dependent. Whereas consistent results were observed after 30 minutes of MNase digestion, extended treatment for 4 h or 24 h resulted in a reduction of median mononucleosomal DNA fragment size to 125 bp and 94 bp, respectively (Supplementary Fig. S1D). For both HCT116 and CAMA-1 cell line conditioned media, MNase digestion for less than 30 minutes resulted in smaller proportions of mononucleosomes (Figure 1D, Supplementary Fig. S1E), with gradual shortening of the mononucleosome-associated fragments over time. Of note, CAMA-1 had more natural mononucleosomes in media before MNase digestion as compared to HCT116, which is possibly related to mechanisms of cell death or intracellular nuclease activity^29^. Taken together, these results demonstrate the activity of the MNase as a nuclease capable of shearing unprotected DNA that slows predictably when encountering the histone core complex within media cf-chromatin^17^.

### Genome-wide media cf-chromatin profiles reflect cell line-specific chromosomal variation

Following demonstration of robust production of nucleosomes from conditioned media across preclinical models, we sought to identify cell type-specific features that could be revealed through genomic profiling. Following MNase treatment, we purified cfDNA from both CAMA-1 and HCT116 and conducted low-pass WGS (1.1X median coverage). This cell-free MNase-Seq (cfMNase-Seq) procedure revealed broad genome-wide concordance between digestion times for a particular cell type but clear distinctions between cell types (Figure 2A, Supplementary Fig. S2A). Interestingly, nucleosomal coverage in the undigested condition was strongly correlated with the various digested conditions (Spearman correlation, CAMA-1: r=0.94 to 0.95, HCT116 r=0.81 to 0.85), indicating that MNase produces physiologic mono-nucleosomes in the media without apparent bias.

**Figure 2.**
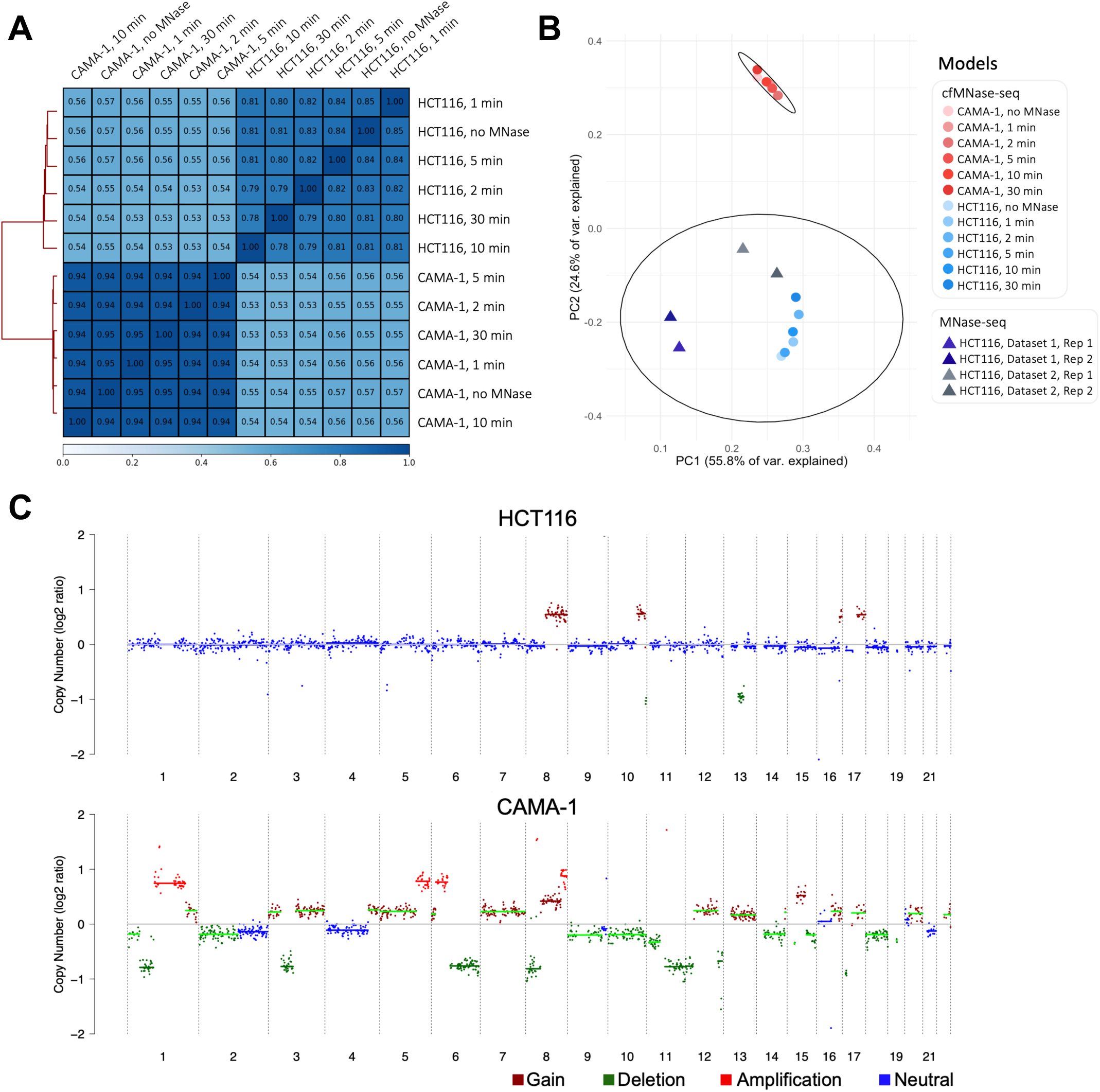
Genome-wide media cf-chromatin profiles reflect cell line-specific chromosomal variation. **A)** Genome-wide Spearman correlation of cfMNase-Seq data across various digestion times for the cell lines HCT116 (n=6) and CAMA-1 (n=6). The correlations were produced over 10,000 bp bins genome-wide. A visual representation of the 10,000 bp bin size is shown in Supplementary Fig. S2A. **B)** PCA of cfMNase-Seq samples across various MNase digestion times (n=12) and samples from two external HCT116 MNase-Seq datasets (n=4). The PCA was performed over 10,000 bp bins genome-wide and clusters were identified using k-means clustering. A silhouette score was calculated to validate clustering; cluster 1 (top) has a silhouette score of 0.98, and cluster 2 (bottom) presents a silhouette score of 0.88. **C)** Copy number analysis with cfMNase-Seq data using ichorCNA. cfMNase-Seq copy number profiles for CAMA-1 compared to WGS, and additional MNase digestion times for HCT116 and CAMA-1 are shown in Supplementary Fig. S2B, S2C, and S2D.

**Figure 3.**
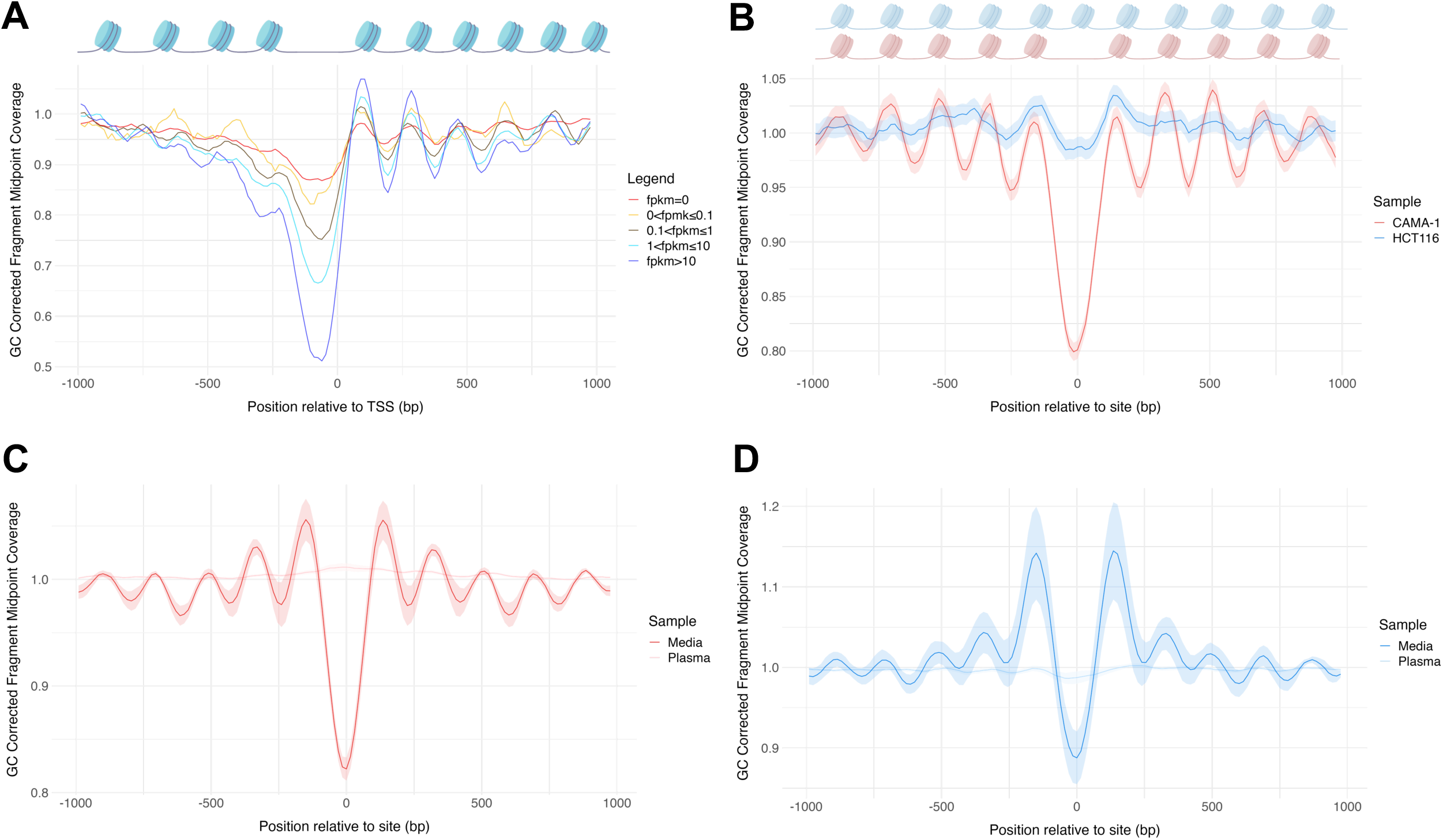
Nucleosome profiles in media cf-chromatin reflect gene expression and chromatin accessibility patterns. **A)** Analysis of the association between cfMNase-Seq coverage and gene expression around the TSS. Composite cfMNase-Seq coverage profiles at TSSs falling within five FPKM levels shown for CAMA-1 (30-minute digestion). Coverage is shown as average GC-corrected fragment midpoint coverage. The complementary analysis for HCT116, information on the FPKM subsets, and coverage profiles across MNase digestion times for HCT116 and CAMA-1 are shown in Supplementary Fig. S3. **B)** Composite cfMNase-Seq coverage profiles (mean ± 95% CI) at 81,292 sites with enriched chromatin accessibility for CAMA-1, shown for CAMA-1 and HCT116 (30-minute digestion). The complementary results for HCT116 enriched open chromatin sites, cfMNase-Seq coverage across all MNase digestions times, and corresponding metrics are shown in Supplementary Fig. S4E-K. **C)** Composite CAMA-1 cfMNase-Seq (n=6) and breast cancer patient plasma WGS (n=54) coverage profiles (mean ± 95% CI) are shown at 215,978 ATAC-Seq open chromatin sites from independent breast cancer patient samples within TCGA. **D)** Composite HCT116 cfMNase-Seq (n=6) and colorectal cancer patient plasma WGS (n=27) coverage profiles (mean ± 95% CI) are shown at 122,971 ATAC-Seq open chromatin sites from independent colorectal cancer patient samples within TCGA. Composite site analysis for all individual samples and the corresponding metrics for C) and D) are in Supplementary Fig. S5.

Next, to confirm that the simulated cf-chromatin was representative of nuclear chromatin organization, we compared CAMA-1 and HCT116 cfMNase-Seq profiles with HCT116 nuclear MNase-Seq profiles obtained from two independent studies^75,76^. Principal component analysis (PCA) of genome-wide nucleosome coverage demonstrated clear distinctions between CAMA-1 and HCT116 cfMNase-Seq profiles (silhouette scores, top cluster=0.98, bottom cluster=0.88), whereas HCT116 nuclear MNase-Seq profiles were more closely related to cfMNase-Seq profiles from HCT116 than CAMA-1 (Figure 2B). This confirms that genome-wide nucleosome profiles revealed by cfMNase-Seq share similarities to the nuclear MNase-Seq signal of the same cell line. As plasma WGS can be used to measure cancer-specific chromosomal structural variants such as copy number aberrations, we suspected that the genome-wide nucleosome coverage revealed by cfMNase-Seq could also be used for this purpose. Therefore, we applied ichorCNA, commonly used for plasma WGS copy number analysis^51^, to cfMNase-Seq profiles from HCT116 and CAMA-1 (Figure 2C). For HCT116, copy number gains were evident for chromosomes 8, 10, 16, and 17, consistent with previous reports^77^. Likewise, for CAMA-1, copy number aberrations detected within cf-chromatin were largely concordant with those from nuclear WGS (Supplementary Fig. S2B)^78^. Furthermore, these copy number profiles were consistent across MNase digestion times (Supplementary Fig. S2C, S2D). Altogether, these results indicate that cfMNase-Seq data reflects cancer-specific copy number profiles and can be analyzed with existing plasma WGS tools.

### Media cf-chromatin sequencing profiles at transcription start sites reflect transcriptional activity

Previous studies on plasma WGS have demonstrated that sequencing coverage of cf-chromatin can be used to deduce nucleosome occupancy maps that, in turn, correlate with cellular transcription profiles^14,18–20^. Specifically, at nucleosome-depleted regions (NDRs) of the genome, which tend to permit binding of transcriptional machinery, there is a reduction in sequencing coverage due to the digestion of unprotected DNA by plasma nucleases. Since nucleosome-protected fragments are over-represented in both plasma WGS and cfMNase-Seq, we asked whether cfMNase-Seq coverage also reflects associations with gene expression.

First, we obtained RNA-Seq data for HCT116 and CAMA-1 and analyzed matched cfMNase-Seq coverage surrounding transcription start sites (TSSs) (Figure 3A, Supplementary Fig. S3A-D). As expected, genes with high expression (e.g., >10 FPKM) displayed a prominent decrease in coverage at the TSS and large oscillations in coverage of flanking regions. This is consistent with strong positioning of nucleosomes upstream and downstream of NDRs^18–20^. In contrast, genes with low expression did not show a pronounced decrease in coverage at the TSS and had weaker oscillations in coverage of flanking regions. The correlation between gene expression and nucleosome positioning was also evident through quantitative metrics, namely mean coverage, central coverage, and amplitude for CAMA-1 (Pearson correlation, mean coverage: r=-0.63, p=2×10^-04^; central coverage: r=-0.88, p=1.4×10^-10^; amplitude: r=0.84, p=5.7×10^-09^), and HCT116 (Pearson correlation, mean coverage: r=0.13, p=0.5; central coverage: r=-0.56, p=0.0012; amplitude: r=0.79, p=1.7×10^-07^). Although strong associations were observed between gene expression and TSS coverage metrics across varying digestion times (Supplementary Fig. S3C-F), the differences became more pronounced as digestion time increased (Supplementary Fig. S3C-F). Overall, these results illustrate how cf-chromatin profiles from preclinical models can reflect transcriptional activity by revealing nuclease-resistant nucleosome-bound regions.

### Media cf-chromatin sequencing profiles beyond transcription start sites reveal cell-specific nucleosome organization

Having demonstrated the ability of cf-chromatin sequencing profiles to reflect nucleosome organization associated with transcriptional activity, we next sought to leverage these profiles to reveal novel cancer-specific chromatin accessibility features distal to TSSs. First, we confirmed that in the vicinity of the TSS, the genes with depletion of cfMNase-Seq coverage corresponded to those with high ATAC-Seq signal in the corresponding cells, indicating that MNase is degrading accessible TSSs (Supplementary Fig. S4A).

Next, we explored cfMNase-Seq profiles at genome-wide chromatin accessibility variants that could distinguish between cell types^79^ (Supplementary Fig. S4B). Using ATAC-Seq data for each cell line, we performed a differential chromatin accessibility analysis and identified regions with enriched accessibility in CAMA-1 and HCT116, respectively (Supplementary Fig. S4B, S4C). Interestingly, these chromatin variants were predominantly found within introns and distal intergenic regions as opposed to the TSS (Supplementary Fig. S4D). For regions with increased accessibility in CAMA-1, we observed a notable decrease in CAMA-1 cfMNase-Seq coverage centered around the chromatin variants with prominent oscillations upstream and downstream, representing an NDR and strongly positioned flanking nucleosomes (Figure 3B). In contrast, the HCT116 cfMNase-Seq signal remained fairly consistent across the CAMA-1 chromatin variants (Figure 3B). As expected, the opposite association was observed for regions with enriched chromatin accessibility in HCT116 (Supplementary Fig. S4E). Among sites with enriched chromatin accessibility in CAMA-1, distinct cfMNase-Seq coverage profiles between CAMA-1 and HCT116 were maintained across all digestion times (Supplementary Fig. S4F), as reflected by quantitative coverage metrics (Supplementary Fig. S4G). Notably, these coverage metrics were also significantly different between CAMA-1 and HCT116 cfMNase-Seq samples (Welch Two Sample t-test, amplitude: p=7.9×10^-05^, mean coverage: p=0.0093; Wilcoxon rank-sum test, central coverage: p=0.0022) (Supplementary Fig. S4H). However, at regions with enriched chromatin accessibility in HCT116, the HCT116 cfMNase-Seq coverage profiles varied notably across digestion times (Supplementary Fig. S4I), with the expected NDR coverage decrease appearing only with longer digestion times (Supplementary Fig. S4J). Among the coverage metrics, solely amplitude displayed a significant difference between CAMA-1 and HCT116 cfMNase-Seq samples (Welch Two Sample t-test, mean coverage: p=0.077, central coverage: p=0.46, amplitude: p=0.0032) (Supplementary Fig. S4K). Taken together, these results indicate that cfMNase-Seq reflects nucleosome organization, chromatin accessibility, and molecular phenotypes that can be used to distinguish cell types.

### Pure tumor-derived media cf-chromatin reflects stronger nucleosome positioning than patient plasma WGS

Lastly, we compared CAMA-1 and HCT116 cfMNase-Seq profiles with plasma WGS profiles from breast and colorectal cancer patients^57^. Using ATAC-Seq consensus peaks from TCGA breast and colorectal cancer tissue to identify independent open chromatin regions^56^, we evaluated cfMNase-Seq and plasma WGS coverage for each respective cancer type. While cfMNase-Seq coverage dropped and reflected strongly positioned surrounding nucleosomes at open chromatin regions from the corresponding cancer type (Figure 3C, 3D and Supplementary Fig. S5A-B), previously published plasma WGS profiles displayed little change in these same regions. Specifically, plasma WGS profiles from breast cancer patients (n=54) had no apparent drop in coverage at the breast cancer open chromatin regions, and the plasma WGS profiles from colorectal cancer patients (n=27) had only a minor dip in coverage at the colorectal cancer open chromatin regions (Figure 3C, 3D and Supplementary Fig. S5C-D). Of note, the colorectal cancer plasma WGS samples contained a higher tumor fraction than the breast cancer samples (Wilcoxon rank sum test, p=0.0067). Nucleosome positioning within open chromatin regions from the respective cancer type was significantly stronger in media cf-chromatin compared with plasma for both breast cancer (Wilcoxon rank sum test, mean coverage: p=0.00012, central coverage: p=6.91×10^-05^, amplitude: p=6.91×10^-05^) and colorectal cancer (Wilcoxon rank sum test, mean coverage: p=0.024, central coverage p=1.81×10^-06^, amplitude: p=1.81×10^-06^). Altogether, these results indicate that pure tumor-derived cf-chromatin sequencing profiles from breast and colorectal cancer conditioned media reflect chromatin accessibility of the corresponding cancer type better than patient plasma WGS.

### H3K4me3 profiling from media cf-chromatin reflects nuclear profiles and cell-specific biology

As nucleosome profiling indicated that cf-chromatin from tissue culture conditioned media maintains the epigenetic state of the cell of origin, we utilized the framework to adapt and develop methods for the comprehensive profiling of genome-wide deposition of histone modifications associated with active and repressive chromatin states^80^ using cfChIP-Seq (Figure 4A; Supplementary Fig. S6A). For this, we chose to study SU-DHL-6, a germinal center B-cell like (GCB) diffuse large B-cell lymphoma (DLBCL) cell line, because of the increasing relevance of epigenetic-targeted therapies in the treatment of lymphomas^81,82^. First, we investigated H3K4me3, a histone modification that reflects active chromatin states at promoter regions responsible for transcriptional regulation^83^ for which proof-of-concept studies have demonstrated the ability of cfChIP-Seq to measure its genome-wide deposition within plasma^21^. Within SU-DHL-6 conditioned media cf-chromatin, H3K4me3 was sharply positioned around TSSs (Figure 4B). This signal was reproduced between replicates and across cf-chromatin input levels (Supplementary Fig. S6B), and peaks called by MACS2 were highly concordant (Figure 4C). Ninety-seven percent of peaks identified by H3K4me3 cfChIP-Seq were also observed within SU-DHL-6 nuclear chromatin from ENCODE^69^ (Figure 4D).

**Figure 4.**
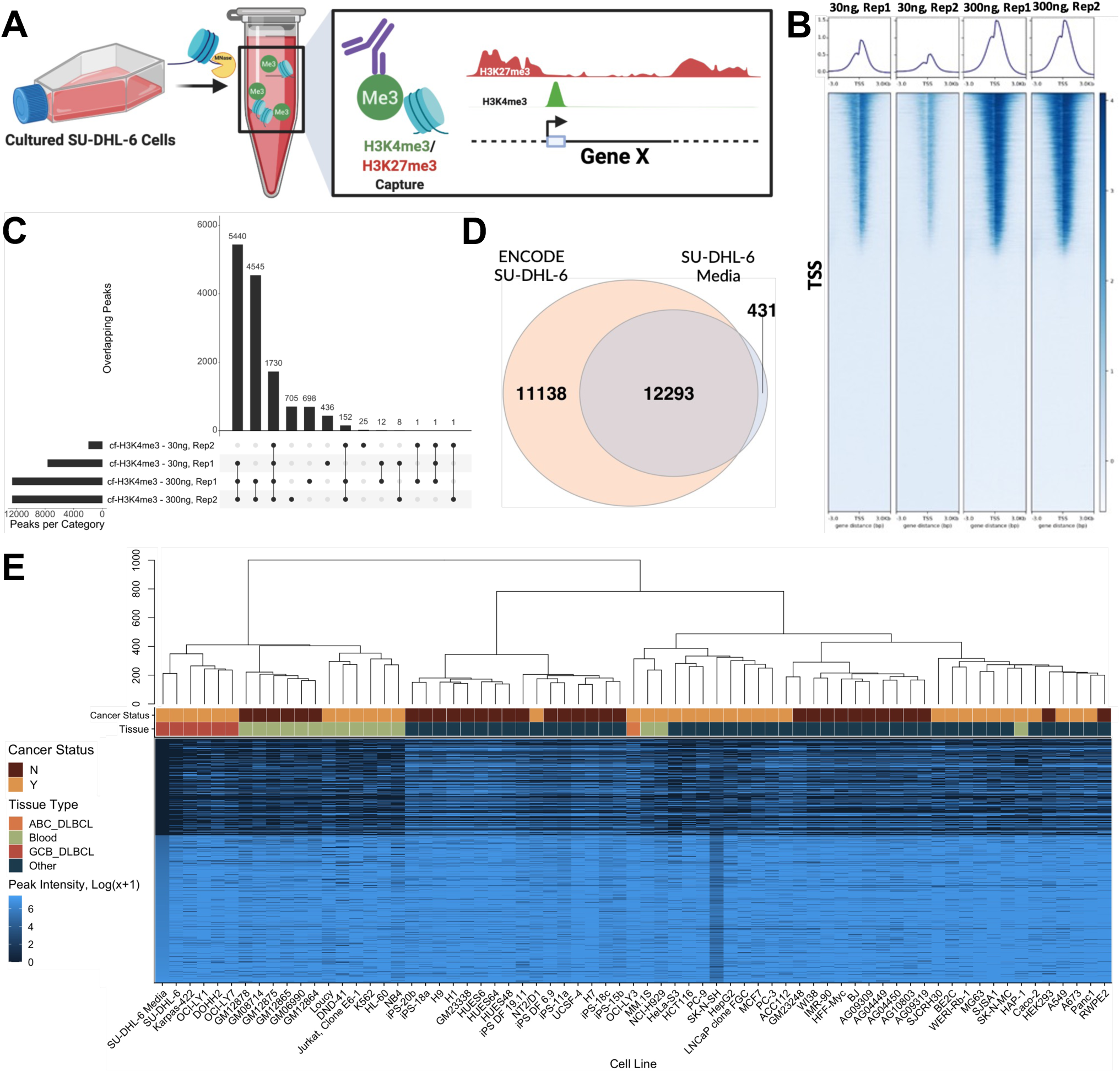
H3K4me3 profiling from media cf-chromatin reflects cell line profiles and biology. **A)** Schematic summarizing methods for MNase treatment of cf-chromatin and cfChIP-Seq for H3K4me3 and H3K27me3 from SU-DHL-6 conditioned media. **B)** H3K4me3 cfChIP-Seq coverage 3 kb around global TSSs, across biological replicates of 30ng (low) and 300ng (high) ChIP input concentrations. Subsampling to the same read depth was conducted before analysis. **C)** UpSet plot of the overlap in H3K4me3 cfChIP-Seq MACS2-called peaks across replicate sets. Peak number for 30ng, Rep2 was low due to low coverage. **D)** Venn diagram representing the overlap in peaks between a cf-H3K4me3 profile (300ng, Rep1) and ENCODE H3K4me3 nuclear chromatin profile from the ENCODE database. **E)** Unsupervised hierarchical clustering of H3K4me3 peak signal intensity (log(x+1) transformed) from SU-DHL-6 media cf-chromatin, SU-DHL-6 chromatin (ENCODE), and other lymphomas (GCB and ABC DLBCLs), other blood-derived or adjacent cell types, and other non-blood cell types (e.g., pancreatic and prostate cancer, and induced pluripotent stem cells). Each row represents the sum of peak intensities at a TSS, and rows were sorted by SU-DHL-6 media H3K4me3 peak intensity.

Having validated the ability to profile H3K4me3 in cf-chromatin from our tissue culture system, we evaluated whether the H3K4me3 cfChIP-Seq profiles were biologically representative of the cell line of interest, along with other similar cell types. We compared H3K4me3 cfChIP-Seq peaks from SU-DHL-6 to H3K4me3 nuclear ChIP-Seq peaks from a spectrum of cell types, including SU-DHL-6, other DLBCL cell lines (GCB and activated B-cell [ABC] subtypes), other hematopoietic cell types, and a mix of differentiated and undifferentiated non-hematopoietic cell types (Figure 4E). Hierarchical clustering of peak intensities at TSSs revealed the similarity of GCB DLBCL profiles, while OCI-LY3, an ABC DLBCL, clustered separately. Cell types from other non-hematopoietic cell types clustered further away from the lymphoma profiles. We also obtained similar results when peaks were mapped to functional pathways prior to clustering (Supplementary Fig. S6C). Together, these results illustrate that H3K4me3 genome-wide deposition within cf-chromatin reflects cell-specific biological states.

### H3K27me3 from media cf-chromatin reflects repressive chromatin states within SU-DHL-6 cells

While H3K4me3 cfChIP-Seq represents a promising source of liquid biopsy biomarkers, we observed only modest variation across cell types, indicating that H3K4me3 alone does not entirely capture the complex epigenetic landscape of transcriptional regulation. We therefore asked whether integration of H3K27me3 cfChIP-Seq profiles could provide complementary signals. H3K27me3 is a heterochromatin mark that spans broad repressive domains associated with treatment response in a variety of cancers^84–86^, and may provide a new source of untapped liquid biopsy biomarkers. Genome-wide distributions of H3K27me3 within SU-DHL-6 cf-chromatin were strongly correlated between replicates and across cf-chromatin input levels (Spearman r=0.96-0.99), and moderately correlated with SU-DHL-6 nuclear chromatin H3K27me3 but not IgG ChIP-Seq from ENCODE^69^ (Figure 5A, Supplementary Fig. S6D). As anticipated, H3K27me3 and RNA-Seq^44^ profiles from SU-DHL-6 cells were negatively correlated, with high RNA expression accompanied by depletion of H3K27me3 from both cf-chromatin and nuclear chromatin. Interestingly, this negative correlation was even stronger for H3K27me3 from cf-chromatin (Spearman correlation, r=-0.47 to −0.50) than from nuclear chromatin (Spearman correlation, r=-0.36 to −0.42), potentially due to reduced background signal from cf-chromatin (Supplementary Fig. S6D).

**Figure 5.**
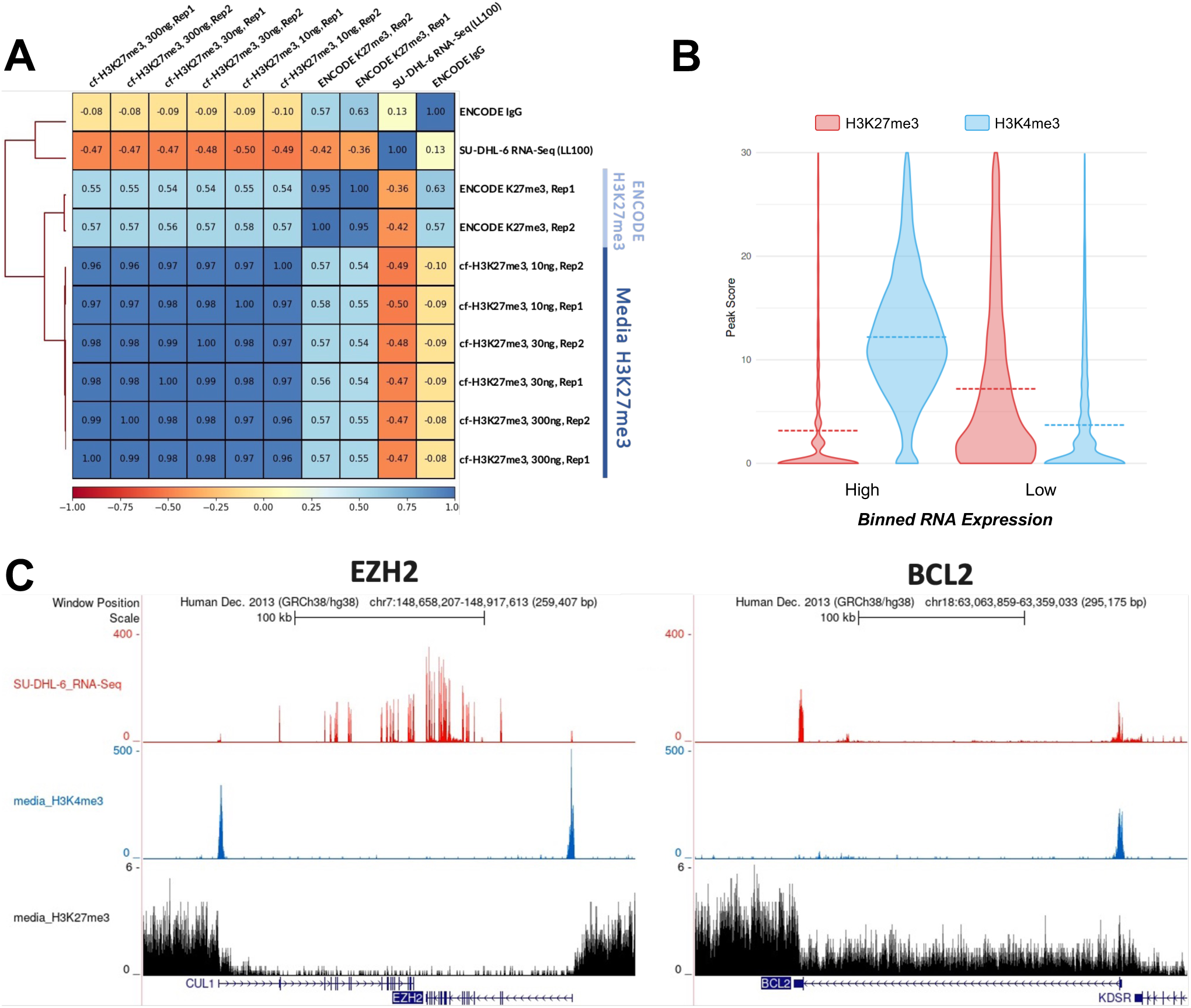
H3K27me3 from media cf-chromatin reflects repressive chromatin states within SU-DHL-6 cells. **A)** Spearman correlation between SU-DHL-6 media cf-H3K27me3 profiles (10, 30, and 300 ng cfDNA input and technical replicates), ENCODE SU-DHL-6 chromatin profiles (biological replicates), SU-DHL-6 RNA-Seq (LL100), and IgG control (ENCODE). **B)** Violin plots depicting SU-DHL-6 media histone modification scores separated by median RNA-Seq expression over 19,802 protein-coding TSSs. Violin plot median values are shown with dashed lines. p-values (H3K4me3, ≤0.0001; H3K27me3, ≤0.0001) were calculated using a two-tailed t-test. **C)** Visualization of SU-DHL-6 RNA-Seq, cf-H3K4me3, and cf-H3K27me3 signal over *EZH2* and *BCL2*, primary drivers of and prognostic factors in DLBCL, respectively.

The vast majority (94%) of peaks from SU-DHL-6 H3K27me3 cfChIP-Seq profiles did not overlap with those from H3K4me3 cfChIP-Seq (Supplementary Fig. S6E). Indeed, consistent with their known antagonistic roles in gene transcriptional regulation^87,88^, these two marks had opposite associations with transcriptional activity in SU-DHL-6 (Figure 5B). Visualization of cfChIP-Seq and RNA-Seq profiles across *EZH2* and *BCL2*, representative genes with established functions in DLBCL^89,90^, revealed depletion of H3K27me3 over the gene bodies and strong enrichment of H3K4me3 at the promoters (Figure 5C). Overall, these results demonstrate that H3K27me3 from media cf-chromatin reflects expected distributions and characteristics of nuclear H3K27me3.

### Active and repressive histone modifications within cf-chromatin predict nuclear chromatin states

In demonstrating that H3K27me3 in media cf-chromatin reflects repressive states, we also observed a subset of regions bivalently marked by both H3K27me3 and H3K4me3 (Supplementary Fig. S5E). Among bivalently marked genomic locations within cf-chromatin, the transcriptional activity of flanking genes varied widely with median levels between those of genes marked by only H3K27me3 or H3K4me3 (Figure 6A). An example of this is *TP53INP1* (Supplementary Fig. S6F), the expression of which has been shown to differentiate lymphomas with *EZH2* mutations and *BCL2* translocations from other lymphoma subtypes^91^. Such bivalency – revealed only through integrated profiling of multiple histone modifications within cf-chromatin – could represent a poised chromatin state compatible with timely adjustments upon developmental cues during tumorigenesis^92^.

**Figure 6.**
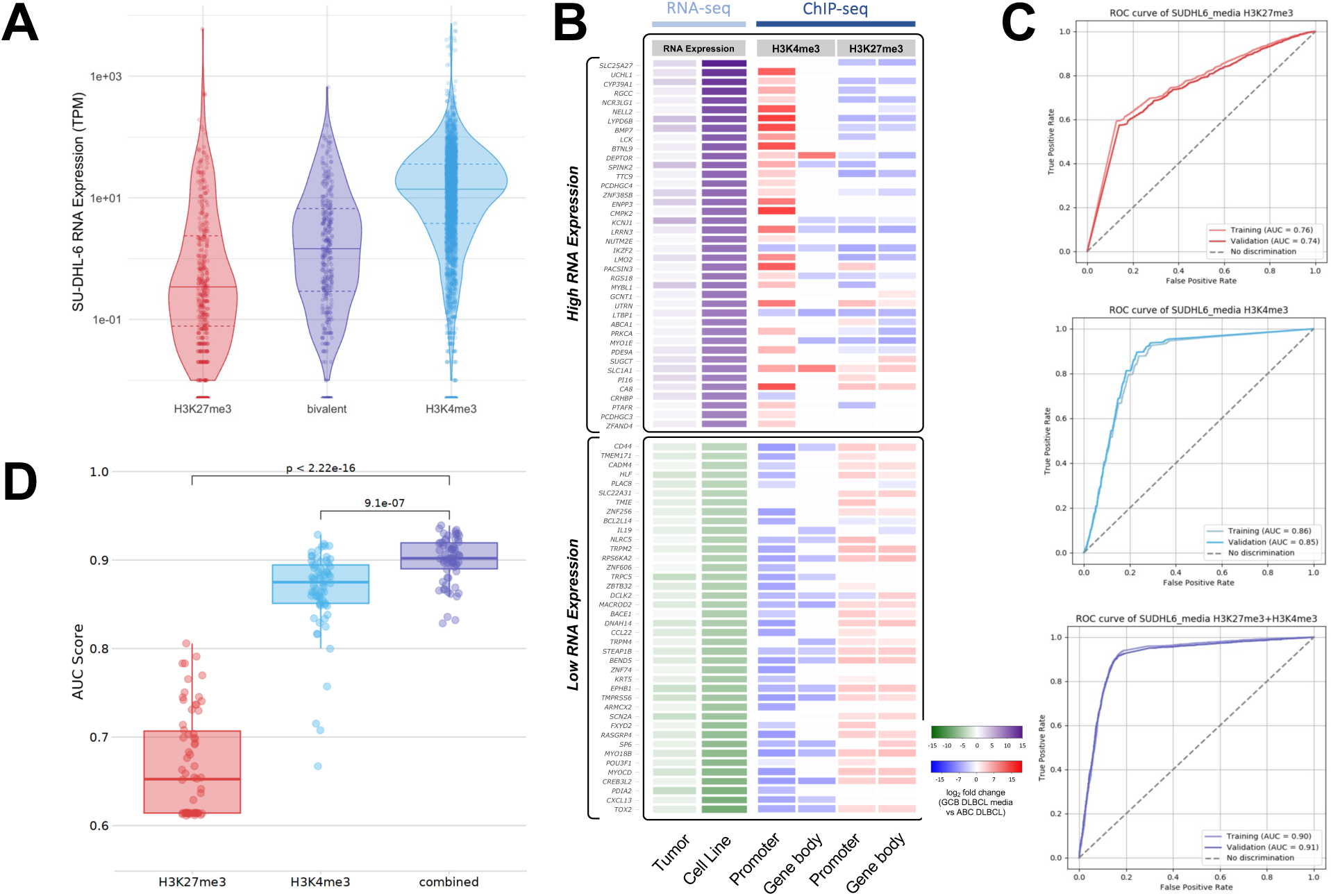
Active and repressive histone modifications within cf-chromatin predict nuclear chromatin states. **A)** Violin plots depicting SU-DHL-6 RNA-Seq expression at genes marked with either promoter H3K27me3 only, H3K4me3 only, or both modifications (i.e., bivalent promoters). Peak groups were identified by quantifying the proportion of overlapping peaks from SU-DHL-6 media H3K4me3 and H3K27me3 profiles (Supplementary Fig. S4B). Only peaks within 3 kb of the TSS of the nearest gene were used in the analysis. Median RNA expression for each group is represented by a solid line, with the interquartile ranges marked by dotted lines. Welch’s t-test: H3K4me3 vs. H3K27me3, p=1.87×10^-7^; H3K4me3 vs. bivalent, p=1.19×10^-11^; H3K27me3 vs. bivalent, p=0.915. **B)** Heatmap of log2 fold change in 60 genes upregulated and downregulated in GCB DLBCL and ABC DLBCL subtypes. SU-DHL-6 (GCB subtype) RNA expression was compared to OCI-LY3 (ABC subtype) along with matching tumor subtypes (left column). We performed pair-wise average ranking between the cell and tumour data based on p-value to select the most significant differentially expressed genes. SU-DHL-6 K27me3 and H3K4me3 cfChIP-Seq data were compared relative to OCI-LY3 ChIP-Seq data. Differential peak intensity for each histone modification is displayed for both upstream promoter or gene body regions. **C)** SU-DHL-6 media receiver operating characteristics (ROC) for training and validation sets for H3K4me3, H3K27me3, and combined histone modification from logistic regression. 10-fold cross-validation was performed using summed histone modification scores over the TSS and gene body of 19,802 protein-coding genes. Model performance was evaluated using the area under the curve (AUC). **D)** Box plots denoting AUC scores of 56 cell types from Roadmap Epigenomics Consortium, 9 cell types from ENCODE, and SU-DHL-6 media (highlighted with a black outline). Paired t-test: combined-H3K4me3 p=9.1×10^-7^; combined-H3K27me3 p=2.2×10^-22^.

We demonstrated previously that SU-DHL-6 (GCB DLBCL), and OCI-LY3 (ABC DLBCL) exhibited unique genome-wide deposition of H3K4me3 within cf-chromatin (Figure 4E). To further elucidate the molecular differences between these two subtypes, we analyzed differential gene expression in the cell line data and patient GCB and ABC DLBCL tumor samples^93^ (Figure 6B). Using genes with differential expression in both cell line and patient samples, we then assessed differential signal intensity from histone modification in the resulting regions. We observed that differentially upregulated genes in SU-DHL-6 relative to OCI-LY3 were enriched for H3K4me3, and diminished for H3K27me3 at their TSS and gene body. The opposite trends in histone modification patterns were observed for genes with lower expression in SU-DHL-6, demonstrating phenotypic characteristics that differentiate the cell lines reflected in cf-chromatin.

Having shown that the combination of active and repressive histone modifications within cf-chromatin may be complementary in reflecting chromatin state, we next assessed the ability of SU-DHL-6 H3K4me3 and H3K27me3 cfChIP-Seq profiles to predict transcriptional activity. We adapted a binary classifier model from Frasca et al.^68^ to predict mRNA expression from histone modifications. SU-DHL-6 cf-chromatin state profiles exhibited robust model performance for each modification and achieved a higher area under the receiver operating characteristic curve (AUC) when both H3K4me3 and H3K27me3 cf-ChIP-Seq profiles were integrated into the model (Figure 6C). Next, to evaluate the performance of H3K27me3 and H3K4me3 to predict transcriptional activity across multiple cell types, we incorporated paired nuclear chromatin histone modification and RNA-Seq data from Roadmap Epigenomics Mapping Consortium (56 cell types) and ENCODE (9 cell types). While AUC values were higher for H3K4me3 than for H3K27me3, the highest AUC values were achieved when incorporating both histone modifications (Figure 6D). Taken together, these results show feasibility of predicting gene expression state using a minimal set of histone modifications from cf-chromatin.

## DISCUSSION

This is the first study dedicated to the simulation of nucleosomal cf-chromatin distributions in plasma using pure tumor-derived cf-chromatin from conditioned tissue culture media. Distinct from cfDNA, cf-chromatin reflects the association with other components including the epigenetically modified histone core complex, the combination of which governs chromatin states and associated DNA-templated processes. Here, we demonstrate an experimental framework using a nuclease treatment to generate nucleosome-sized cf-chromatin consistently in both 2-D and 3-D culture systems. An advantage of simulated cf-chromatin in tissue culture media is the absence of interfering signals from hematopoietic cells and other tissues. Herein, we used simulated cf-chromatin to discover pure tumor-derived molecular features through comprehensive genome-wide profiling of nucleosome positioning, active histone modifications, and for the first time in a liquid biopsy context, a repressive histone modification. As a renewable source of cf-chromatin, our simulated cf-chromatin framework is ideal for liquid biopsy methods development and validation prior to testing with patient samples. Moreover, in this work, we assessed our novel cf-chromatin datatypes to nuclear chromatin profiles from the cell of origin to reinforce the biological representation of this system. Compared to nuclear chromatin for liquid biopsy biomarker development, simulated cf-chromatin is a comparable model to plasma cf-chromatin, permits easy and repeated sampling, and allows for preprocessing (e.g., cfDNA isolation) and analysis (e.g., short read sequencing) techniques that are compatible with liquid biopsy technologies. Altogether, we propose that simulated cf-chromatin and the profiling techniques showcased herein can be used for biomarker discovery for further interrogation in patient samples.

In this work, we ascertained chromatin states within cf-chromatin by conducting cfMNase-Seq as well as cfChIP-Seq for both H3K4me3 and H3K27me3. Using cfMNase-Seq, we found that highly expressed genes displayed an NDR near their respective TSS and strong positioning of surrounding nucleosomes, which is consistent with WGS performed on plasma cf-chromatin^18–20^. We also expanded beyond a gene-centric view by investigating genome-wide chromatin variants that could identify distinct cfMNase-Seq profiles. Notably, sites with enriched chromatin accessibility in CAMA-1 had a much clearer separation of coverage profiles between CAMA-1 and HCT116 than those with enriched accessibility in HCT116. We found fewer regions of accessible chromatin enriched in HCT116 compared to CAMA-1, which may have limited the coverage analysis and be related to the CpG island methylator phenotype of HCT116^94^. Next, we explored cfMNase-Seq and patient plasma WGS coverage across accessible chromatin sites determined by ATAC-Seq data from patient tumor tissue samples. While regions of accessible chromatin in patient tumor samples corresponded with decreased cfMNase-Seq coverage, these pronounced coverage changes were not observed in patient plasma WGS. This highlights the limitations of plasma WGS in which low fractional abundance of ctDNA dramatically dilutes tumor-derived cf-chromatin features. In contrast, the simulated cf-chromatin analysis framework presented here provides unencumbered access to tumor-derived chromatin accessibility features without background contribution of signal from other cell types. Altogether, these results demonstrate that cfMNase-Seq reflects the chromatin landscape of the cell of origin, and is generalizable to patient tumor chromatin accessibility of the corresponding cancer type, demonstrating stronger nucleosome positioning patterns than patient plasma WGS.

As we leveraged simulated cf-chromatin to develop cfChIP-Seq methods for H3K27me3 profiling, this study marks the initial exploration of cfChIP-Seq profiles for histone modifications associated with both active and repressive chromatin states. As previously observed^21^, the active mark H3K4me3 was sharply positioned at a subset of TSSs within cf-chromatin. The distribution of H3K4me3 within SU-DHL-6 cf-chromatin was most similar to nuclear chromatin ChIP-Seq profiles from SU-DHL-6 and other GCB DLBCL cell lines while being distinct from an ABC DLBCL cell line and non-hematopoietic cell types. Since ABC DLBCLs are phenotypically and genetically different from GCB DLBCLs^95^, we examined differences in cf-chromatin H3K4me3 and H3K27me3. H3K27me3 marks repressive chromatin domains where transcriptional activity should be low, which is consistent with our findings in SU-DHL-6. We observed distinct histone modification patterns within genes that are differentially expressed between GCB and ACB DLBCL^96^ including *LYPD6B*^97^*, TOX2*^98^, and *CD44*^99^. With growing evidence that subtyping lymphoma can improve patient outcomes^100^, cfChIP-Seq may offer new opportunities for clinical utility of epigenomic liquid biopsy biomarkers.

Integrative profiling of both H3K4me3 and H3K27me3 from cf-chromatin may allow for more accurate prediction of chromatin state from the cell of origin. Approximately 6% of H3K27me3 peaks and 9% of H3K4me3 peaks were shared in SU-DHL-6 cf-chromatin. Promoters sharing these modifications marked genes with an intermediate expression phenotype, better reflecting chromatin state. Bivalent domains have also been shown to be an integral part of cellular development and lineage specification^89,101^. Profiling both marks within cf-chromatin could improve the phenotypic characterization of certain cancers that undergo dramatic epigenomic dysregulation, resulting in significant changes in gene expression and chromosome stability^102^. In the case of DLBCL, a cancer linked to epigenomic dysregulation, integrative cfChIP-Seq analysis may provide deeper insights for predictive biomarkers. Furthermore, we found that models incorporating both H3K4me3 and H3K27me3 significantly increased the accuracy of gene expression binary classification. Extending the application of cfChIP-Seq to incorporate a wider range of histone modifications could have the potential to improve biomarker discovery. Overall, we demonstrate that our simulated cf-chromatin framework is applicable to both novel liquid biopsy methods development, and biological exploration for new epigenetic biomarkers.

Despite our efforts to validate the simulation of cf-chromatin from tissue culture and its downstream epigenetic profiling, our work has certain limitations. First, this framework uses MNase, which is derived from bacteria and may not produce the same cfDNA fragmentation features as nucleases in human plasma (e.g., DNase1L3, DNase1)^103^. Second, we focused on three cell lines for most of the epigenomic profiling analyses; future work should confirm the applicability of these techniques in additional models. Third, we utilized low coverage cfMNase-Seq (∼1X), which inherently limited our ability to evaluate nucleosome footprints and gene expression at single-gene resolution that could be explored in deep sequencing datasets. Finally, we selected two histone modifications for cfChIP-Seq profiling, whereas numerous others have been implicated in defining chromatin states relevant to cellular phenotypes^104^; future work could seek to incorporate other histone modifications and even DNA methylation (e.g., cfMeDIP-Seq)^10,11^ into this new simulated cf-chromatin framework.

Moreover, xenograft mouse models have also been used for cf-chromatin biomarker discovery^16,29^ and may better account for physiologic variables affecting cfDNA release and digestion; however, they are labor-intensive and can produce low quantities and fractions of tumor-derived cf-chromatin from small plasma samples. Future work could attempt to overcome these limitations and directly compare our simulated cf-chromatin to that from xenograft models. Areas for further exploration also include leveraging this framework to evaluate epigenetic changes associated with drug treatments, transformation, or differentiation over time with serial cf-chromatin samples.

In summary, we have developed a framework for the simulation of cf-chromatin using conditioned media from cultured 2-D and 3-D models that better emulates cf-chromatin topology in plasma. This framework provides readily accessible cf-chromatin material, enabling liquid biopsy biomarker exploration without the contribution of background signal, which currently burdens exploratory analysis in human plasma samples. We also demonstrate the use of simulated cf-chromatin in liquid biopsy profiling methods development, as we adapted and developed multiple profiling approaches for evaluating various chromatin states, including cfMNase-Seq and cfChIP-Seq for H3K4me3 and H3K27me3. Altogether, our findings showcase the benefit of epigenetic profiling of pure tumor-derived cf-chromatin from preclinical models. We intend for our framework of simulated cf-chromatin and downstream profiling techniques to be applied in preclinical models to produce priors for biomarker discovery that can be tested in limited clinical samples. We anticipate this work will accelerate the development of minimally invasive biomarkers applicable to cancer detection, classification, and treatment response prediction or monitoring, contributing to the future of precision medicine.

## DATA AVAILABILITY

New sequence data generated in this study and associated code to generate figures in this manuscript will be made available at the time of publication. Publicly available datasets utilized in this manuscript are available as follows. The MNase-Seq FASTQ files for HCT116 were downloaded from the European Nucleotide Archive (https://www.ebi.ac.uk/ena/browser/view/SRX2349700 [SRR5022986 and SRR5022987]; https://www.ebi.ac.uk/ena/browser/view/SRX6066529 [SRR9297149]; https://www.ebi.ac.uk/ena/browser/view/SRX6066530 [SRR9297150])^75,76^. The WGS FASTQ files^78^ for CAMA-1 were downloaded (available at 10.5281/zenodo.6760819). The plasma WGS BAM files from treatment-naive breast and colorectal cancer patients were downloaded from the European Genome-phenome Archive at the European Bioinformatics Institute (EGAD00001005339) with agreed use by the User Institution with a data access agreement^57^; the tumor fractions for EGAD00001005339 were obtained from a previous publication^15^ that ran ichorCNA on all samples. The ATAC-Seq FASTQ files for HCT116 were downloaded from ENCODE^69^ (https://www.encodeproject.org/experiments/ENCSR872WGW/). The RNA-Seq FASTQ files for HCT116 were downloaded from ENCODE (https://www.encodeproject.org/experiments/ENCSR698RPL/). The RNA-Seq FASTQ files for SU-DHL-6 ([SRR19938568]^105^, [ERR3003598]^44^) and OCILY3 ([SRR974972]^106^, [SRR17188140]^107^) were downloaded from Sequence Read Archive (SRA) and European Nucleotide Archive (ENA). The RNA-Seq data for 111 DLBCL specimens of formalin-fixed paraffin-embedded tissues were downloaded from NCBI (BioProject: PRJNA622950).

The H3K4me3 and H3K27me3 profiles from assorted cell types used in various analyses were downloaded from the Roadmap Epigenomics Consortium (Supplementary Table S3) and ENCODE database (Supplementary Table S4).

## ABBREVIATIONS

cfDNA: 
cf-chromatin: 
MNase: 
WGS: 
cfChIP-Seq: 
cfMNase-Seq: 
TSSs: 
NDR: 
ATAC-Seq: 
BRCA: 
COAD: 
DLBCL: 
GBC: 
ABC: 

## ACKNOWLEDGEMENTS

We thank Jennifer Cruickshank for providing organoid conditioned media samples and Samah El Ghamrasni for helping set up the RNA-Seq analysis pipeline. We thankfully acknowledge the Princess Margaret Genomics Centre for carrying out the next-generation sequencing and Zhibin Lu (University Health Network High-Performance Computing Centre and Bioinformatics Core) for technical assistance. We also thank members of the Bratman laboratory for helpful discussion and suggestions. The graphical abstract and schematics in Figures 3A, 3B, 4A, S1D, S3A, and S4B were created using BioRender.

## FUNDING

S.V.B. is supported by the Gattuso-Slaight Personalized Cancer Medicine Fund at the Princess Margaret Cancer Centre and the Dr. Mariano Elia Chair in Head & Neck Cancer Research at the University Health Network and the University of Toronto. S.D. is supported by a Canadian Institute of Health Research (CIHR) Fredrick Banting and Charles Best Canada Graduate Doctoral Scholarship (CGS-D, Application #457244). D.W.C is supported by the Canadian Institutes of Health Research and the Princess Margaret Cancer Foundation (DH Gales Family Foundation and the Green Fischer Family Trust and Goldie R. Feldman). S.C.M. is supported by the Cancer Digital Intelligence Spark Award at the Princess Margaret Cancer Centre and the Ontario Graduate Scholarship at the University of Toronto.

## COMPETING INTERESTS

S.V.B. is inventor on patents related to cell-free DNA mutation and methylation analysis technologies that are unrelated to this work and have been licensed to Roche Molecular Diagnostics and Adela, respectively. S.V.B. is a co-founder of, has ownership in, and serves in a leadership role at Adela. M.M.H. is an inventor on a patent application related to cell-free methylation analysis licensed to Adela. D.W.C reports consulting/advisory roles with AstraZeneca, Exact Sciences, Eisai, Gilead, GlaxoSmithKline, Inflex, Inivata, Merck, Novartis, Pfizer, Roche and Saga; research funding to institution from AstraZeneca, Guardant Health, Gilead, GlaxoSmithKline, Inivata, Knight, Merck, Pfizer, and Roche and Patent (US62/675,228) for methods of treating cancers characterized by a high expression level of spindle and kinetochore associated complex subunit 3 (ska3) gene. R.K. reports research funding from Roche and Abbvie. All other authors declare no conflicts.

## REFERENCES

1. Lone, S. N. et al. Liquid biopsy: a step closer to transform diagnosis, prognosis and future of cancer treatments. Mol. Cancer 21, 79 (2022).

2. Lo, Y. M. D., Han, D. S. C., Jiang, P. & Chiu, R. W. K. Epigenetics, fragmentomics, and topology of cell-free DNA in liquid biopsies. Science 372, (2021).

3. Moss, J. et al. Comprehensive human cell-type methylation atlas reveals origins of circulating cell-free DNA in health and disease. Nat. Commun. (2018) doi:10.1038/s41467-018-07466-6.

4. Heitzer, E., Haque, I. S., Roberts, C. E. S. & Speicher, M. R. Current and future perspectives of liquid biopsies in genomics-driven oncology. Nature Reviews Genetics Preprint at 10.1038/s41576-018-0071-5 (2019).

5. Ezeife, D. A. et al. The economic value of liquid biopsy for genomic profiling in advanced non-small cell lung cancer. Ther. Adv. Med. Oncol. 14, 17588359221112696 (2022).

6. Newman, A. M. et al. An ultrasensitive method for quantitating circulating tumor DNA with broad patient coverage. Nat. Med. (2014) doi:10.1038/nm.3519.

7. Zviran, A. et al. Genome-wide cell-free DNA mutational integration enables ultra-sensitive cancer monitoring. Nat. Med. 26, 1114–1124 (2020).

8. Moser, T., Kühberger, S., Lazzeri, I., Vlachos, G. & Heitzer, E. Bridging biological cfDNA features and machine learning approaches. Trends Genet. (2023) doi:10.1016/j.tig.2023.01.004.

9. Im, Y. R., Tsui, D. W. Y., Diaz, L. A., Jr & Wan, J. C. M. Next-Generation Liquid Biopsies: Embracing Data Science in Oncology. Trends Cancer Res. 7, 283–292 (2021).

10. Shen, S. Y., et al. Sensitive tumour detection and classification using plasma cell-free DNA methylomes. Nature Preprint at 10.1038/s41586-018-0703-0 (2018).

11. Burgener, J. M. et al. Tumor-Naïve Multimodal Profiling of Circulating Tumor DNA in Head and Neck Squamous Cell Carcinoma. Clin. Cancer Res. 27, 4230–4244 (2021).

12. Song, C. X. et al. 5-Hydroxymethylcytosine signatures in cell-free DNA provide information about tumor types and stages. Cell Res. (2017) doi:10.1038/cr.2017.106.

13. Vorperian, S. K., Moufarrej, M. N., Tabula Sapiens Consortium & Quake, S. R. Cell types of origin of the cell-free transcriptome. Nat. Biotechnol. (2022) doi:10.1038/s41587-021-01188-9.

14. Snyder, M. W., Kircher, M., Hill, A. J., Daza, R. M. & Shendure, J. Cell-free DNA Comprises an in Vivo Nucleosome Footprint that Informs Its Tissues-Of-Origin. Cell 164, 57–68 (2016).

15. Doebley, A.-L. et al. A framework for clinical cancer subtyping from nucleosome profiling of cell-free DNA. Nat. Commun. 13, 7475 (2022).

16. De Sarkar, N. et al. Nucleosome patterns in circulating tumor DNA reveal transcriptional regulation of advanced prostate cancer phenotypes. bioRxiv 2022.06.21.496879 (2022) doi:10.1101/2022.06.21.496879.

17. Chereji, R. V., Bryson, T. D. & Henikoff, S. Quantitative MNase-seq accurately maps nucleosome occupancy levels. Genome Biol. (2019) doi:10.1186/s13059-019-1815-z.

18. Ulz, P. et al. Inferring expressed genes by whole-genome sequencing of plasma DNA. Nat. Genet. 48, 1273–1278 (2016).

19. Zhu, G. et al. Tissue-specific cell-free DNA degradation quantifies circulating tumor DNA burden. Nat. Commun. 12, 2229 (2021).

20. Esfahani, M. S. et al. Inferring gene expression from cell-free DNA fragmentation profiles. Nat. Biotechnol. 40, 585–597 (2022).

21. Sadeh, R. et al. ChIP-seq of plasma cell-free nucleosomes identifies gene expression programs of the cells of origin. Nat. Biotechnol. (2021) doi:10.1038/s41587-020-00775-6.

22. Fedyuk, V. et al. Multiplexed, single-molecule, epigenetic analysis of plasma-isolated nucleosomes for cancer diagnostics. Nat. Biotechnol. 41, 212–221 (2023).

23. Roadmap Epigenomics Consortium et al. Integrative analysis of 111 reference human epigenomes. Nature 518, 317–330 (2015).

24. Phallen, J. et al. Direct detection of early-stage cancers using circulating tumor DNA. Sci. Transl. Med. (2017) doi:10.1126/scitranslmed.aan2415.

25. Moding, E. J., Nabet, B. Y., Alizadeh, A. A. & Diehn, M. Detecting Liquid Remnants of Solid Tumors: Circulating Tumor DNA Minimal Residual Disease. Cancer Discov. 11, 2968–2986 (2021).

26. Bettegowda, C. et al. Detection of circulating tumor DNA in early- and late-stage human malignancies. Sci. Transl. Med. 6, 224ra24 (2014).

27. Abbosh, C., Birkbak, N. J. & Swanton, C. Early stage NSCLC — challenges to implementing ctDNA-based screening and MRD detection. Nature Reviews Clinical Oncology Preprint at 10.1038/s41571-018-0058-3 (2018).

28. Ungerer, V., Bronkhorst, A. J., Van den Ackerveken, P., Herzog, M. & Holdenrieder, S. Serial profiling of cell-free DNA and nucleosome histone modifications in cell cultures. Sci. Rep. 11, 9460 (2021).

29. Rostami, A. et al. Senescence, Necrosis, and Apoptosis Govern Circulating Cell-free DNA Release Kinetics. Cell Rep. 31, 107830 (2020).

30. Sachs, N. et al. A Living Biobank of Breast Cancer Organoids Captures Disease Heterogeneity. Cell 172, 373–386.e10 (2018).

31. Rago, C. et al. Serial assessment of human tumor burdens in mice by the analysis of circulating DNA. Cancer Res. (2007) doi:10.1158/0008-5472.CAN-07-0605.

32. Rostami, A., Yu, C. & Bratman, S. V. Serial Cell-free DNA Assessments in Preclinical Models. STAR Protocols 1, 100145 (2020).

33. Wang, T. T. et al. High efficiency error suppression for accurate detection of low-frequency variants. Nucleic Acids Res. 47, 1–11 (2019).

34. Corces, M. R. et al. An improved ATAC-seq protocol reduces background and enables interrogation of frozen tissues. Nat. Methods 14, 959–962 (2017).

35. Picelli, S. et al. Tn5 transposase and tagmentation procedures for massively scaled sequencing projects. Genome Res. 24, 2033–2040 (2014).

36. Martin, M. Cutadapt removes adapter sequences from high-throughput sequencing reads. EMBnet.journal 17, 10–12 (2011).

37. Langmead, B. & Salzberg, S. L. Fast gapped-read alignment with Bowtie 2. Nat. Methods 9, 357–359 (2012).

38. Danecek, P. et al. Twelve years of SAMtools and BCFtools. Gigascience 10, (2021).

39. Ramírez, F. et al. deepTools2: a next generation web server for deep-sequencing data analysis. Nucleic Acids Res. 44, W160–5 (2016).

40. Zhang, Y. et al. Model-based analysis of ChIP-Seq (MACS). Genome Biol. 9, R137 (2008).

41. Amemiya, H. M., Kundaje, A. & Boyle, A. P. The ENCODE Blacklist: Identification of Problematic Regions of the Genome. Sci. Rep. 9, 9354 (2019).

42. Dobin, A. et al. STAR: ultrafast universal RNA-seq aligner. Bioinformatics 29, 15– 21 (2013).

43. Li, B. & Dewey, C. N. RSEM: accurate transcript quantification from RNA-Seq data with or without a reference genome. BMC Bioinformatics 12, 323 (2011).

44. Quentmeier, H. et al. The LL-100 panel: 100 cell lines for blood cancer studies. Sci. Rep. (2019) doi:10.1038/s41598-019-44491-x.

45. Kim, D. et al. TopHat2: accurate alignment of transcriptomes in the presence of insertions, deletions and gene fusions. Genome Biol. 14, R36 (2013).

46. Lander, E. S. et al. Initial sequencing and analysis of the human genome. Nature 409, 860–921 (2001).

47. Merchant, N. et al. The iPlant Collaborative: Cyberinfrastructure for Enabling Data to Discovery for the Life Sciences. PLoS Biol. 14, e1002342 (2016).

48. Kent, W. J. et al. The Human Genome Browser at UCSC. Genome Research vol. 12 996–1006 Preprint at 10.1101/gr.229102 (2002).

49. Hadley Wickham. ggplot2: Elegant Graphics for Data Analysis.

50. Ha, G. et al. Integrative analysis of genome-wide loss of heterozygosity and monoallelic expression at nucleotide resolution reveals disrupted pathways in triple-negative breast cancer. Genome Res. 22, 1995–2007 (2012).

51. Adalsteinsson, V. A. et al. Scalable whole-exome sequencing of cell-free DNA reveals high concordance with metastatic tumors. Nat. Commun. 8, 1324 (2017).

52. Stark, R. & Brown, G. DiffBind : Differential binding analysis of ChIP-Seq peak data.

53. Ross-Innes, C. S. et al. Differential oestrogen receptor binding is associated with clinical outcome in breast cancer. Nature 481, 389–393 (2012).

54. Love, M. I., Huber, W. & Anders, S. Moderated estimation of fold change and dispersion for RNA-seq data with DESeq2. Genome Biol. 15, 1–21 (2014).

55. Yu, G., Wang, L.-G. & He, Q.-Y. ChIPseeker: an R/Bioconductor package for ChIP peak annotation, comparison and visualization. Bioinformatics 31, 2382–2383 (2015).

56. Corces, M. R. et al. The chromatin accessibility landscape of primary human cancers. Science 362, eaav1898 (2018).

57. Cristiano, S. et al. Genome-wide cell-free DNA fragmentation in patients with cancer. Nature 570, 385–389 (2019).

58. Karolchik, D. et al. The UCSC Table Browser data retrieval tool. Nucleic Acids Res. 32, D493–6 (2004).

59. Foundation for Statistical Computing, R. R. R: a language and environment for statistical computing. RA Lang Environ Stat Comput.

60. Galili, T. dendextend: an R package for visualizing, adjusting and comparing trees of hierarchical clustering. Bioinformatics 31, 3718–3720 (2015).

61. Patwardhan, M., Wenger, C., Davis, E., and Phanstiel, D. (2022). bedtoolsr: Bedtools Wrapper. R package version 2.30.0-4.

62. Durinck, S., Spellman, P. T., Birney, E. & Huber, W. Mapping identifiers for the integration of genomic datasets with the R/Bioconductor package biomaRt. Nat. Protoc. 4, 1184–1191 (2009).

63. Neuwirth, E. (2022). RColorBrewer: ColorBrewer Palettes. R package version 1.1-3.

64. Mills, B.R. (2022). MetBrewer: Color Palettes Inspired by Works at the Metropolitan Museum of Art. R package version 0.2.0.

65. Welch, R. P. et al. ChIP-Enrich: gene set enrichment testing for ChIP-seq data. Nucleic Acids Res. 42, e105 (2014).

66. Carroll, T. S., Liang, Z., Salama, R., Stark, R. & de Santiago, I. Impact of artifact removal on ChIP quality metrics in ChIP-seq and ChIP-exo data. Front. Genet. 5, 75 (2014).

67. Bioconductor Core Team and Bioconductor Package Maintainer (2021). TxDb.Hsapiens.UCSC.hg38.knownGene: Annotation package for TxDb object(s). R package version 3.14.0.

68. Frasca, F., Matteucci, M., Leone, M., Morelli, M. J. & Masseroli, M. Accurate and highly interpretable prediction of gene expression from histone modifications. BMC Bioinformatics 23, 151 (12/2022).

69. Consortium, E. P. et al. An integrated encyclopedia of DNA elements in the human genome. Nature (2013) doi:10.1038/nature11247.An.

70. Luo, Y. et al. New developments on the Encyclopedia of DNA Elements (ENCODE) data portal. Nucleic Acids Res. 48, D882–D889 (2020).

71. Kagda, M. S. et al. Data navigation on the ENCODE portal. arXiv [q-bio.GN] (2023).

72. Hitz, B. C., et al. The ENCODE Uniform Analysis Pipelines. bioRxiv (2023) doi:10.1101/2023.04.04.535623.

73. Brind’Amour, J. et al. An ultra-low-input native ChIP-seq protocol for genome-wide profiling of rare cell populations. Nat. Commun. (2015) doi:10.1038/ncomms7033.

74. Skene, P. J. & Henikoff, S. An efficient targeted nuclease strategy for high-resolution mapping of DNA binding sites. Elife (2017) doi:10.7554/eLife.21856.

75. Guzman, C. & D’Orso, I. CIPHER: a flexible and extensive workflow platform for integrative next-generation sequencing data analysis and genomic regulatory element prediction. BMC Bioinformatics 18, 363 (2017).

76. Bacon, C. W. et al. KAP1 Is a Chromatin Reader that Couples Steps of RNA Polymerase II Transcription to Sustain Oncogenic Programs. Mol. Cell 78, 1133– 1151.e14 (2020).

77. Passerini, V. et al. The presence of extra chromosomes leads to genomic instability. Nat. Commun. 7, 10754 (2016).

78. Soria-Bretones, I. et al. The spindle assembly checkpoint is a therapeutic vulnerability of CDK4/6 inhibitor-resistant ER+ breast cancer with mitotic aberrations. Sci. Adv. 8, eabq4293 (2022).

79. Grillo, G. & Lupien, M. Cancer-associated chromatin variants uncover the oncogenic role of transposable elements. Curr. Opin. Genet. Dev. 74, 101911 (2022).

80. Zhao, Z. & Shilatifard, A. Epigenetic modifications of histones in cancer. Genome Biol. 20, 245 (2019).

81. Morschhauser, F. et al. Tazemetostat for patients with relapsed or refractory follicular lymphoma: an open-label, single-arm, multicentre, phase 2 trial. Lancet Oncol. 21, 1433–1442 (2020).

82. He, M. Y. & Kridel, R. Treatment resistance in diffuse large B-cell lymphoma. Leukemia (2021) doi:10.1038/s41375-021-01285-3.

83. Wang, H. et al. H3K4me3 regulates RNA polymerase II promoter-proximal pause-release. Nature 615, 339–348 (2023).

84. Marsolier, J. et al. H3K27me3 conditions chemotolerance in triple-negative breast cancer. Nat. Genet. 54, 459–468 (2022).

85. Deblois, G. et al. Epigenetic switch-induced viral mimicry evasion in chemotherapy-resistant breast cancer. Cancer Discov. 10, 1312–1329 (2020).

86. Gardner, E. E. et al. Chemosensitive Relapse in Small Cell Lung Cancer Proceeds through an EZH2-SLFN11 Axis. Cancer Cell (2017) doi:10.1016/j.ccell.2017.01.006.

87. Barth, T. K. & Imhof, A. Fast signals and slow marks: the dynamics of histone modifications. Trends Biochem. Sci. 35, 618–626 (2010).

88. Yu, J.-R., Lee, C.-H., Oksuz, O., Stafford, J. M. & Reinberg, D. PRC2 is high maintenance. Genes Dev. 33, 903–935 (2019).

89. Béguelin, W. et al. EZH2 is required for germinal center formation and somatic EZH2 mutations promote lymphoid transformation. Cancer Cell 23, 677–692 (2013).

90. Iqbal, J. et al. BCL2 expression is a prognostic marker for the activated B-cell--like type of diffuse large B-cell lymphoma. J. Clin. Oncol. 24, 961–968 (2006).

91. Xu-Monette, Z. Y. et al. Genetic subtyping and phenotypic characterization of the immune microenvironment and MYC/BCL2 double expression reveal heterogeneity in diffuse large B-cell lymphoma. Clin. Cancer Res. 28, 972–983 (2022).

92. Macrae, T. A., Fothergill-Robinson, J. & Ramalho-Santos, M. Regulation, functions and transmission of bivalent chromatin during mammalian development. Nat. Rev. Mol. Cell Biol. 24, 6–26 (2023).

93. Yan, W.-H. et al. Cell-of-Origin Subtyping of Diffuse Large B-Cell Lymphoma by Using a qPCR-based Gene Expression Assay on Formalin-Fixed Paraffin-Embedded Tissues. Front. Oncol. 10, 803 (2020).

94. Berg, K. C. G. et al. Multi-omics of 34 colorectal cancer cell lines - a resource for biomedical studies. Mol. Cancer 16, 116 (2017).

95. Schmitz, R. et al. Genetics and Pathogenesis of Diffuse Large B-Cell Lymphoma. N. Engl. J. Med. 378, 1396–1407 (2018).

96. Shipp, M. A. et al. Diffuse large B-cell lymphoma outcome prediction by gene-expression profiling and supervised machine learning. Nat. Med. 8, 68–74 (2002).

97. Hodkinson, B. P. et al. Biomarkers of response to ibrutinib plus nivolumab in relapsed diffuse large B-cell lymphoma, follicular lymphoma, or Richter’s transformation. Transl. Oncol. 14, 100977 (2021).

98. Zhou, J. et al. Super-enhancer-driven TOX2 mediates oncogenesis in Natural Killer/T Cell Lymphoma. Mol. Cancer 22, 69 (2023).

99. Tzankov, A. et al. Prognostic significance of CD44 expression in diffuse large B cell lymphoma of activated and germinal centre B cell-like types: a tissue microarray analysis of 90 cases. J. Clin. Pathol. 56, 747–752 (2003).

100. Nowakowski, G. S. & Czuczman, M. S. ABC, GCB, and Double-Hit Diffuse Large B-Cell Lymphoma: Does Subtype Make a Difference in Therapy Selection? Am Soc Clin Oncol Educ Book e449–57 (2015).

101. Madani Tonekaboni, S. A., Haibe-Kains, B. & Lupien, M. Large organized chromatin lysine domains help distinguish primitive from differentiated cell populations. Nat. Commun. 12, 499 (2021).

102. Baylin, S. B. & Jones, P. A. Epigenetic Determinants of Cancer. Cold Spring Harb. Perspect. Biol. 8, (2016).

103. Han, D. S. C. & Lo, Y. M. D. The Nexus of cfDNA and Nuclease Biology. Trends Genet. 37, 758–770 (2021).

104. Millán-Zambrano, G., Burton, A., Bannister, A. J. & Schneider, R. Histone post-translational modifications - cause and consequence of genome function. Nat. Rev. Genet. 23, 563–580 (2022).

105. Bal, E. et al. Super-enhancer hypermutation alters oncogene expression in B cell lymphoma. Nature 607, 808–815 (2022).

106. Hardee, J. et al. STAT3 targets suggest mechanisms of aggressive tumorigenesis in diffuse large B-cell lymphoma. G3 3, 2173–2185 (2013).

107. Kwon, D. et al. Targeting Refractory Mantle Cell Lymphoma for Imaging and Therapy Using C-X-C Chemokine Receptor Type 4 Radioligands. Clin. Cancer Res. 28, 1628–1639 (2022).

108. Shah, R. N. et al. Examining the Roles of H3K4 Methylation States with Systematically Characterized Antibodies. Mol. Cell (2018) doi:10.1016/j.molcel.2018.08.015.

109. Nassar, L. R. et al. The UCSC Genome Browser database: 2023 update. Nucleic Acids Res. 51, D1188–D1195 (2023).

