## Supplemental Figures and Tables for "Simulating cell-free chromatin using preclinical models for cancer-specific biomarker discovery"

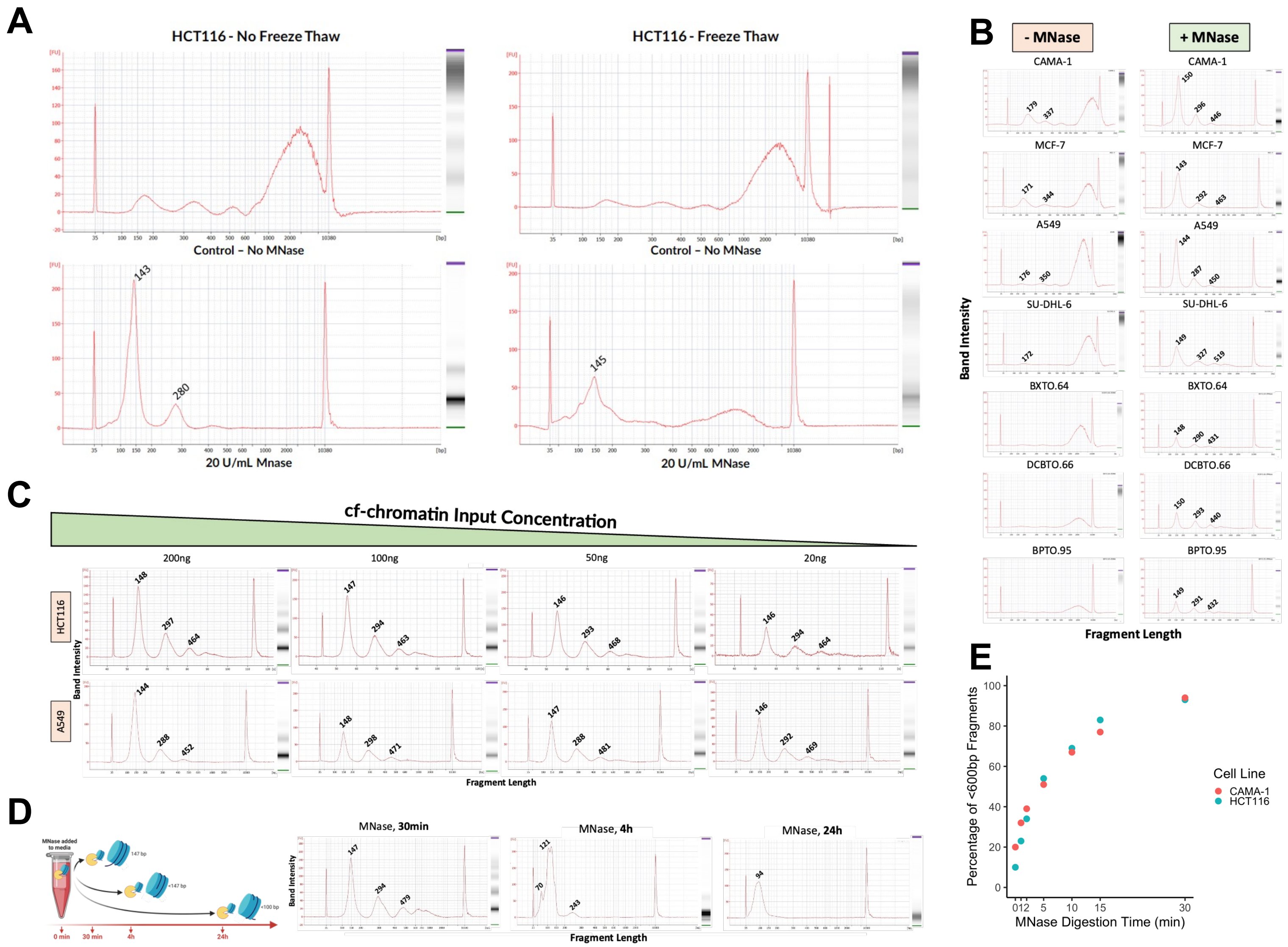

A
HCT116 - No Freeze Thaw
HCT116 - Freeze Thaw
Control – No MNase
20 U/mL Mnase
20 U/mL MNase
143
280
145
B
- MNase
+ MNase
CAMA-1
MCF-7
A549
SU-DHL-6
BXTO.64
DCBTO.66
BPTO.95
Band Intensity
Fragment Length
C
cf-chromatin Input Concentration
200ng
100ng
50ng
20ng
HCT116
A549
Band Intensity
Fragment Length
D
MNase added to media
0 min
30 min
4h
24h
MNase, 30min
MNase, 4h
MNase, 24h
Band Intensity
Fragment Length
E
Percentage of <600bp Fragments
MNase Digestion Time (min)
Cell Line
CAMA-1
HCT116

**Supplementary Figure 1. Methodological considerations for MNase treatment of media cf-chromatin. A)** Results of Agilent BioAnalyzer analysis are shown. HCT116 culture media cf-chromatin was treated with MNase before and after media freeze-thaw compared to untreated conditions (left and right, respectively). **B)** Comparison of MNase vs no MNase conditions for all cell lines and breast cancer organoid models are shown. **C)** Different concentrations of HCT116 media cf-chromatin (ranging from 20-200 ng) were treated with MNase in 1 mL volumes, respectively. Bioanalyzer analysis after cfDNA purification was performed. **D)** HCT116 media was treated with MNase and the reaction halted after 30 minutes, four hours, and 24 h. After 30 minutes, mono-, di- and tri-nucleosomes are present. As digestion continues, oligonucleosome fragments are degraded, and MNase continues to digest mono-nucleosomes within the nucleosome core. **E)** Progressive nucleosome production over time is consistent across HCT116 and CAMA-1 media, with a higher rate of nucleosomes produced at earlier time points.

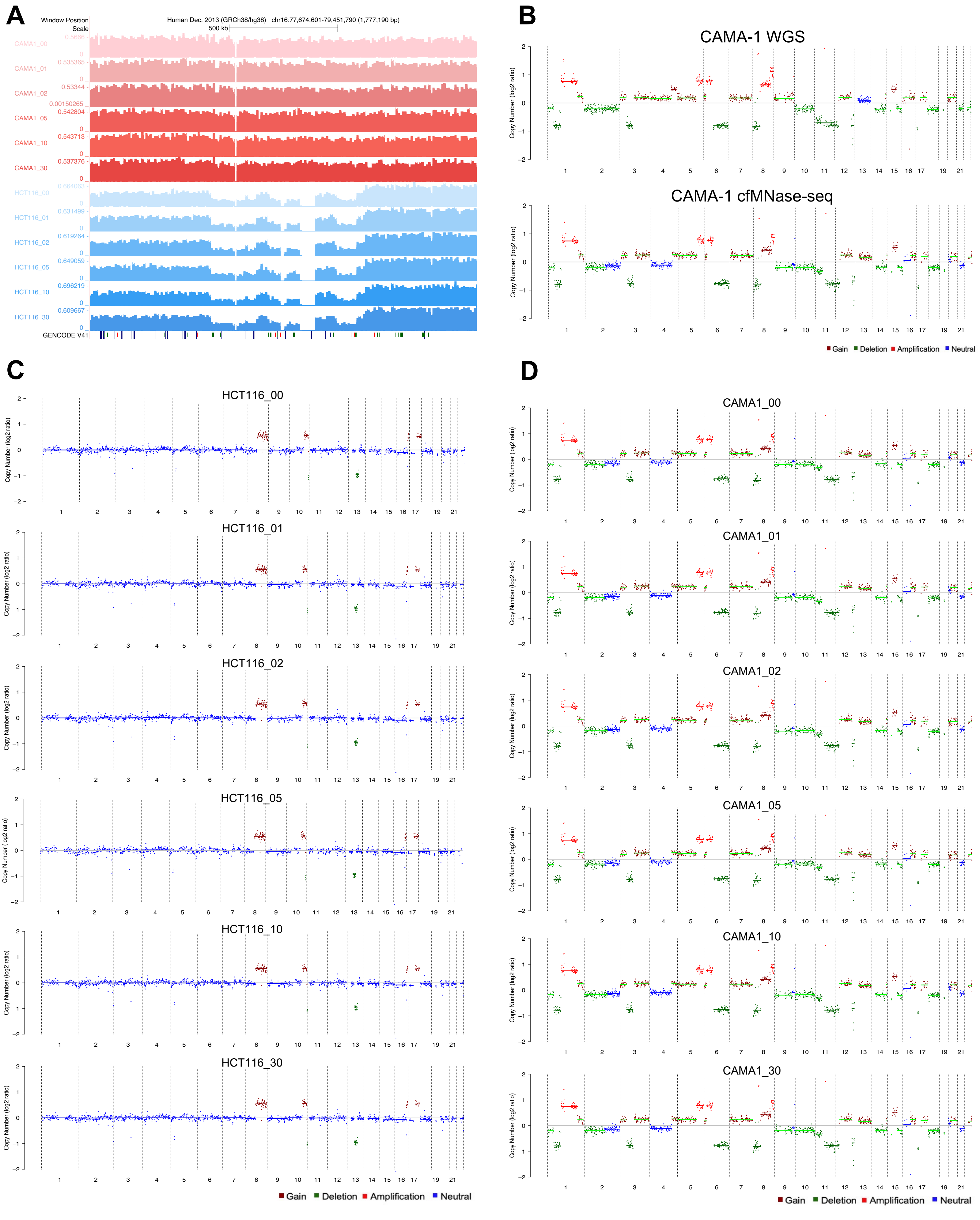
A
Window Position
Scale
Human Dec. 2013 (GRCh38/hg38) chr16:77,674,601-79,451,790 (1,777,190 bp)
500 kb
CAMA1_00
CAMA1_01
CAMA1_02
CAMA1_05
CAMA1_10
CAMA1_30
HCT116_00
HCT116_01
HCT116_02
HCT116_05
HCT116_10
HCT116_30
GENCODE V41
B
CAMA-1 WGS
CAMA-1 cfMNase-seq
Copy Number (log2 ratio)
Gain Deletion Amplification Neutral
C
HCT116_00
HCT116_01
HCT116_02
HCT116_05
HCT116_10
HCT116_30
D
CAMA1_00
CAMA1_01
CAMA1_02
CAMA1_05
CAMA1_10
CAMA1_30

**Supplementary Figure 2. Genome-wide cfMNase-Seq profiles are consistent across different digestion times and concordant with nuclear signal. A)** A representation of the 10,000bp bin size resolution through a UCSC Genome Browser view. This bin size was used for genome-wide comparisons across cfMNase-Seq samples. **B)** Copy number analysis comparison to WGS data and cfMNase-Seq (30-minute digestion) for CAMA-1. **C)** Copy number analysis using ichorCNA with cfMNase-Seq data across various digestion times (no MNase to 30-minute digestion, from top to bottom) for HCT116. **D)** Copy number analysis using ichorCNA with cfMNase-Seq data across various digestion times (no MNase to 30-minute digestion, from top to bottom) for CAMA-1.

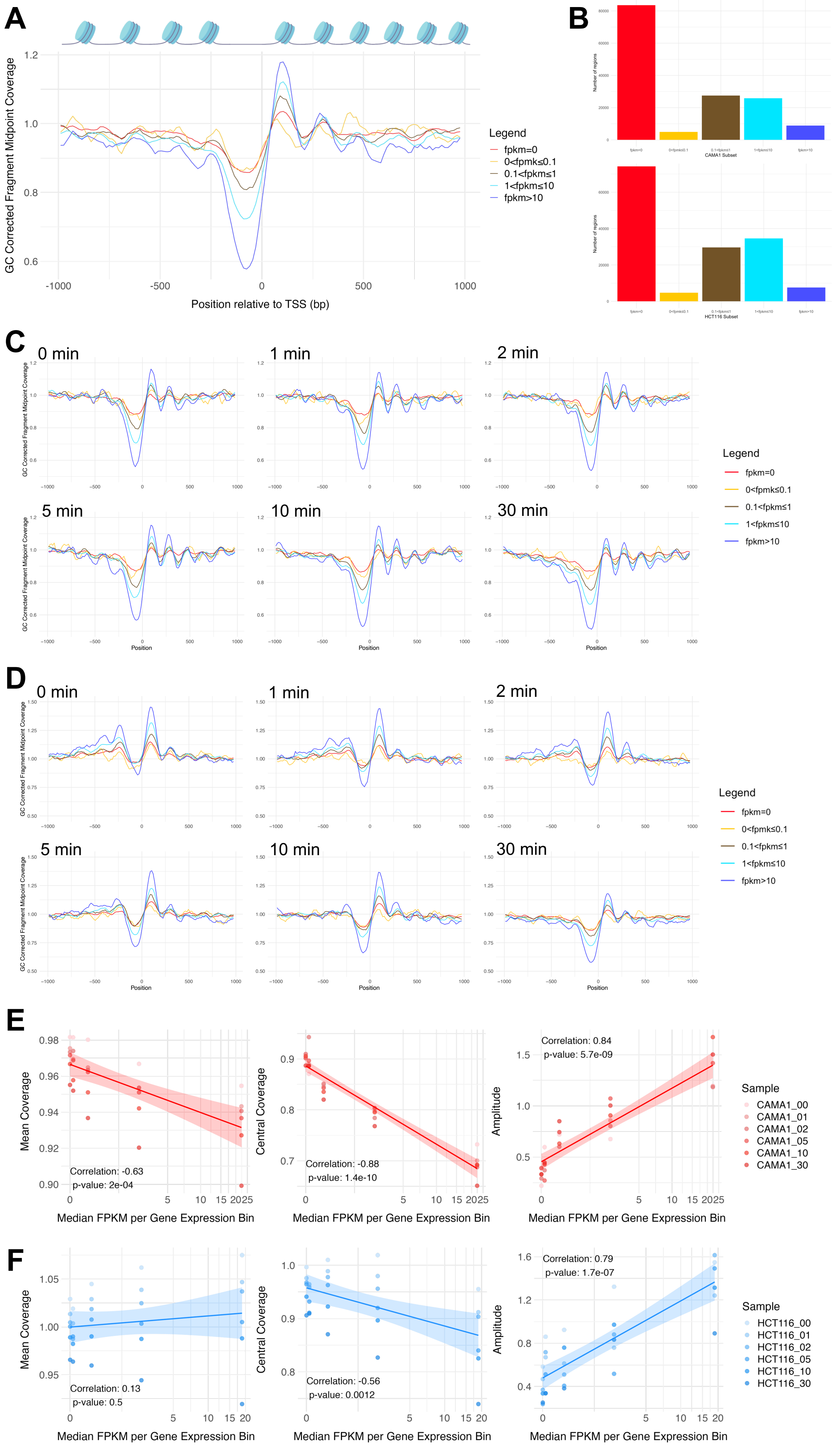

A
GC Corrected Fragment Midpoint Coverage
Position relative to TSS (bp)
Legend
fpkm=0
0<fpmk≤0.1
0.1<fpkm≤1
1<fpkm≤10
fpkm>10
B
Number of regions
CAMA1 Subset
HCT116 Subset
C
0 min
1 min
2 min
5 min
10 min
30 min
Position
D
E
Mean Coverage
Correlation: -0.63
p-value: 2e-04
Central Coverage
Correlation: -0.88
p-value: 1.4e-10
Amplitude
Correlation: 0.84
p-value: 5.7e-09
Median FPKM per Gene Expression Bin
Sample
CAMA1_00
CAMA1_01
CAMA1_02
CAMA1_05
CAMA1_10
CAMA1_30
F
Correlation: 0.13
p-value: 0.5
Correlation: -0.56
p-value: 0.0012
Correlation: 0.79
p-value: 1.7e-07
HCT116_00
HCT116_01
HCT116_02
HCT116_05
HCT116_10
HCT116_30

**Supplementary Figure 3. cfMNase-Seq coverage is associated with gene expression around TSSs. A)** Composite cfMNase-Seq coverage profiles at TSSs falling within five FPKM levels shown for HCT116 (30-minute digestion). Coverage is shown as the average GC-corrected fragment midpoint coverage. **B)** A bar graph representing the number of TSSs that fall into the five FPKM subsets (FPKM=0, 0<FPKM≤0.1, 0.1<FPKM≤1, 1<FPKM≤10, FPKM>10) for CAMA-1 (top) and HCT116 (bottom). **C)** cfMNase-Seq coverage profiles around the TSS for various gene expression levels and across MNase digestion times for CAMA-1. **D)** cfMNase-Seq coverage profiles around the TSS for various gene expression levels and across MNase digestion times for HCT116. **E)** Quantitative metrics from cfMNase-Seq coverage profiles across different gene expression levels (Pearson correlation: mean coverage r=-0.63, p=2x10$^{-04}$; central coverage r=-0.88, p=1.4x10$^{-10}$; amplitude r=0.84, p=5.7x10$^{-09}$), shown for CAMA-1 samples of different digestion times. The quantitative metrics are mean coverage (i.e., average across the entire window), central coverage (i.e., the average coverage of 60 bp centered around the TSS), and amplitude (i.e., periodicity reflecting the strength of positioning and occupancy of nucleosomes). **F)** Quantitative metrics from cfMNase-Seq coverage profiles across different gene expression levels (Pearson correlation: mean coverage r=0.13, p=0.5; central coverage r=-0.56, p=0.0012; amplitude r=0.79, p=1.7x10$^{-07}$), shown for HCT116 samples of different digestion times.

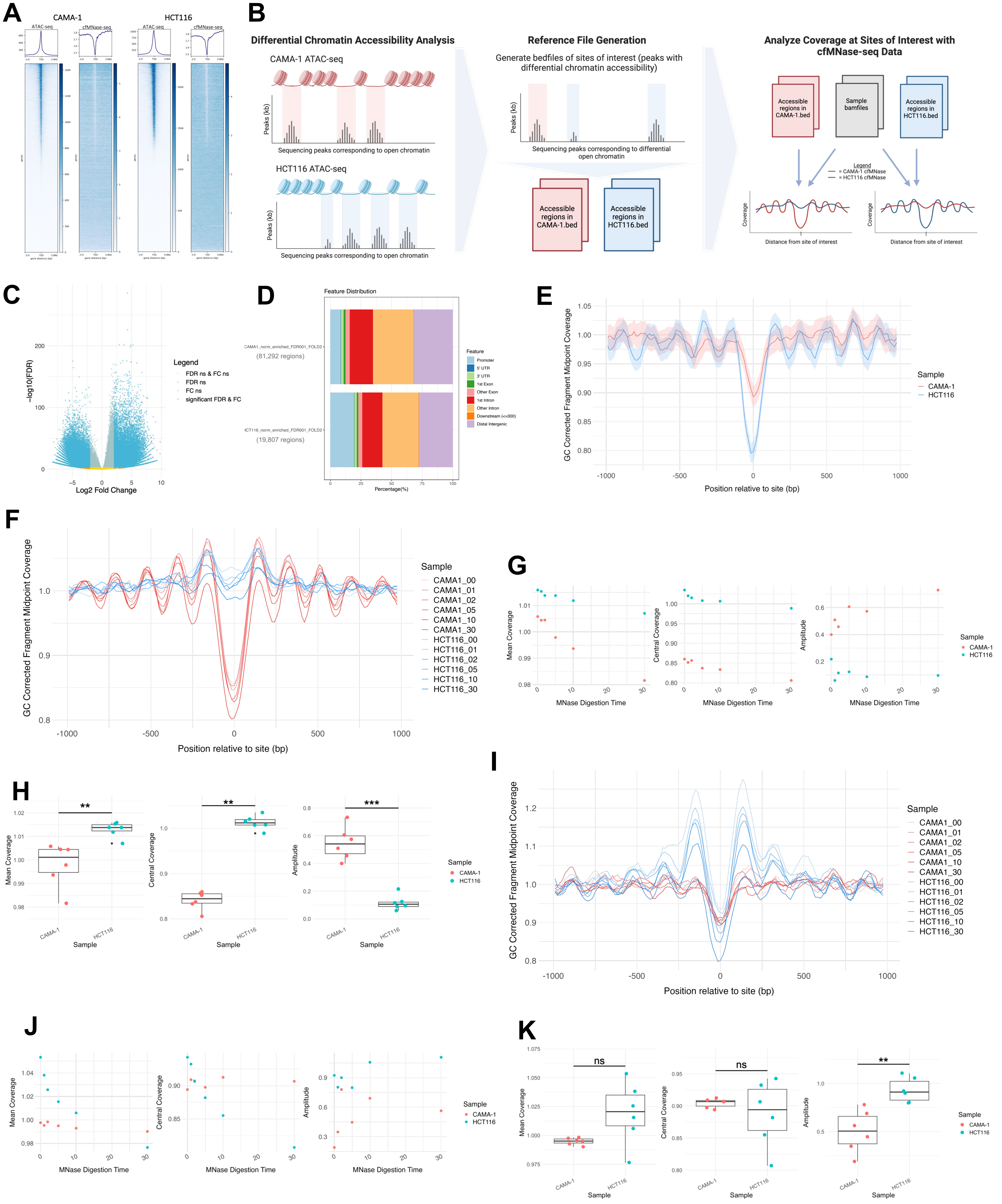

A
CAMA-1
HCT116
ATAC-seq
cfMNase-seq
B
Differential Chromatin Accessibility Analysis
CAMA-1 ATAC-seq
Peaks (kb)
Sequencing peaks corresponding to open chromatin
HCT116 ATAC-seq
Reference File Generation
Generate bedfiles of sites of interest (peaks with differential chromatin accessibility)
Sequencing peaks corresponding to differential open chromatin
Accessible regions in CAMA-1.bed
Accessible regions in HCT116.bed
Analyze Coverage at Sites of Interest with cfMNase-seq Data
Sample bamfiles
Legend
= CAMA-1 cfMNase
= HCT116 cfMNase
Coverage
Distance from site of interest
C
−log10(FDR)
Log2 Fold Change
Legend
FDR ns & FC ns
FDR ns
FC ns
significant FDR & FC
D
Feature Distribution
CAMA1_norm_enriched_FDR001_FOLD2
(81,292 regions)
HCT116_norm_enriched_FDR001_FOLD2
(19,807 regions)
Feature
Promoter
5' UTR
3' UTR
1st Exon
Other Exon
1st Intron
Other Intron
Downstream (<=300)
Distal Intergenic
Percentage(%)
E
GC Corrected Fragment Midpoint Coverage
Position relative to site (bp)
Sample
CAMA-1
HCT116
F
GC Corrected Fragment Midpoint Coverage
Position relative to site (bp)
Sample
CAMA1_00
CAMA1_01
CAMA1_02
CAMA1_05
CAMA1_10
CAMA1_30
HCT116_00
HCT116_01
HCT116_02
HCT116_05
HCT116_10
HCT116_30
G
Mean Coverage
Central Coverage
Amplitude
MNase Digestion Time
Sample
CAMA-1
HCT116
H
**
**
***
Mean Coverage
Central Coverage
Amplitude
Sample
CAMA-1
HCT116
I
GC Corrected Fragment Midpoint Coverage
Position relative to site (bp)
Sample
J
Mean Coverage
Central Coverage
Amplitude
MNase Digestion Time
Sample
K
ns
ns
**
Mean Coverage
Central Coverage
Amplitude
Sample

**Supplementary Figure 4. cfMNase-Seq coverage is associated with chromatin accessibility. A)** Analysis of the association of chromatin accessibility and cfMNase-Seq coverage around the TSS. Rows in the heatmap are individual genes, sorted by the ATAC-Seq signal of the corresponding cell line. **B)** Schematic of the differential chromatin accessibility analysis between CAMA-1 and HCT116 as regions of interest for cfMNase-Seq coverage analysis. Differential sites were identified with CAMA-1 and HCT116 ATAC-Seq peaks using DESeq2 within diffbind. Sites with a log2 fold change greater than two or less than negative two and a false discovery rate less than 0.01 were counted as differential and were used to assess cfMNase-Seq coverage. **C)** Volcano plot of differential sites considered for cfMNase-Seq coverage analysis. Sites with a log2 fold change greater than two or less than negative two and a false discovery rate less than 0.01 were considered differential (shown in blue); all others were not considered for cfMNase-Seq coverage evaluation (FDR=false discovery rate, FC=fold change, ns=not significant). **D)** Distribution of genomic features for differential sites where CAMA1_norm_enriched_FDR001_FOLD denotes sites with enriched accessibility in CAMA-1 (81,292 sites), and HCT116_norm_enriched_FDR001_FOLD represents sites with enriched accessibility in HCT116 (19,807 sites). **E)** Composite cfMNase-Seq coverage profiles (mean ± 95% CI) at 19,807 sites with enriched chromatin accessibility for HCT116, shown for CAMA-1 and HCT116 (30-minute digestion). **F)** Composite cfMNase-Seq coverage profiles at 81,292 sites with enriched chromatin accessibility for CAMA-1, shown for CAMA-1 and HCT116 across all digestion times. **G)** Quantitative metrics (mean coverage, central coverage, and amplitude, reflecting the average coverage across the entire window, 60 bp centered at the zero position, and periodicity reflecting the strength of nucleosome positioning) from cfMNase-Seq coverage profiles at sites with enriched chromatin accessibility for CAMA-1, shown for CAMA-1 and HCT116 across different digestion times. **H)** Comparison of CAMA-1 and HCT116 nucleosome positioning metrics at sites with enriched chromatin accessibility in CAMA-1(Welch Two Sample t-test, amplitude: $p=7.9 \times 10^{-05}$, mean coverage: p=0.0093; Wilcoxon rank-sum test, central coverage: p=0.0022). **I)** Composite cfMNase-Seq coverage profiles at 19,807 sites with enriched chromatin accessibility for HCT116, shown for CAMA-1 and HCT116 across all digestion times. **J)** Quantitative metrics from cfMNase-Seq coverage profiles at sites with enriched chromatin accessibility for HCT116, shown for CAMA-1 and HCT116 across different digestion times. **K)** Comparison of CAMA-1 and HCT116 nucleosome positioning metrics at sites with enriched chromatin accessibility in HCT116 (Welch Two Sample t-test: mean coverage p=0.077; central coverage p=0.46, amplitude p=0.0032).

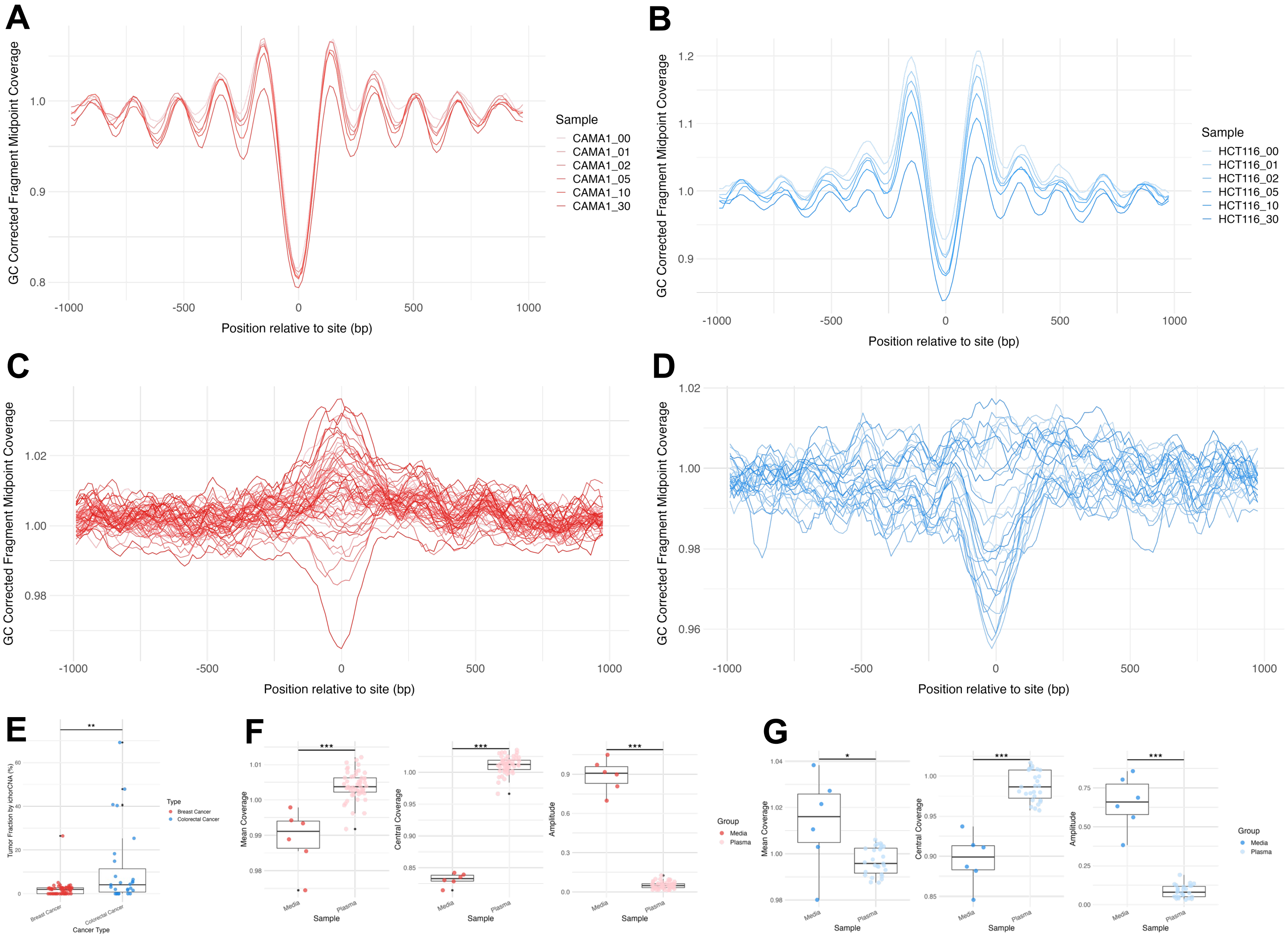

A
GC Corrected Fragment Midpoint Coverage
Position relative to site (bp)
Sample
CAMA1_00
CAMA1_01
CAMA1_02
CAMA1_05
CAMA1_10
CAMA1_30
B
HCT116_00
HCT116_01
HCT116_02
HCT116_05
HCT116_10
HCT116_30
C
D
E
Tumor Fraction by ichorCNA (%)
Type
Breast Cancer
Colorectal Cancer
Cancer Type
F
Mean Coverage
Central Coverage
Amplitude
Sample
Media
Plasma
Group
G

**Supplementary Figure 5. cfMNase-Seq reveals stronger nucleosome positioning patterns than patient plasma WGS at open chromatin regions from patient tissue. A)** Composite cfMNase-Seq coverage profiles at open chromatin regions from breast cancer patient tumors ATAC-Seq consensus peaks from TCGA (215,978 sites) shown for all digestion times of CAMA-1 (n=6). **B)** Composite cfMNase-Seq coverage profiles at open chromatin regions from colorectal cancer patient tumors ATAC-Seq consensus peaks from TCGA (122,971 sites) shown for all digestion times of HCT116 (n=6). **C)** Composite coverage profiles at open chromatin regions from breast cancer patient tumors ATAC-Seq consensus peaks from TCGA (215,978 sites) shown for all breast cancer patient plasma WGS samples (n=54). **D)** Composite coverage profiles at open chromatin regions from colorectal cancer patient tumors ATAC-Seq consensus peaks from TCGA (122,971 sites) shown for all colorectal cancer patient plasma WGS samples (n=27). **E)** Comparision of plasma WGS samples tumor fractions for breast cancer (median=2.00, IQR=0-2.75) and colorectal cancer (median=4.10, IQR=0.78-11.42), as determined by ichorCNA (Wilcoxon rank sum test, p=0.0067). **F)** Comparison of breast cancer plasma and media cf-chromatin nucleosome positioning metrics at open chromatin regions from breast cancer patient tumors ATAC-Seq consensus peaks from TCGA (215,978 sites) (Wilcoxon rank sum test: mean coverage p=0.00012; central coverage $p=6.91x10^{-05}$, amplitude $p=6.91x10^{-05}$). **G)** Comparison of colorectal cancer plasma and media cf-chromatin nucleosome positioning metrics at open chromatin regions from colorectal cancer patient tumors ATAC-Seq consensus peaks from TCGA (122,971 sites) (Wilcoxon rank sum test: mean coverage p=0.024; central coverage $p=1.81x10^{-06}$, amplitude $p=1.81x10^{-06}$).

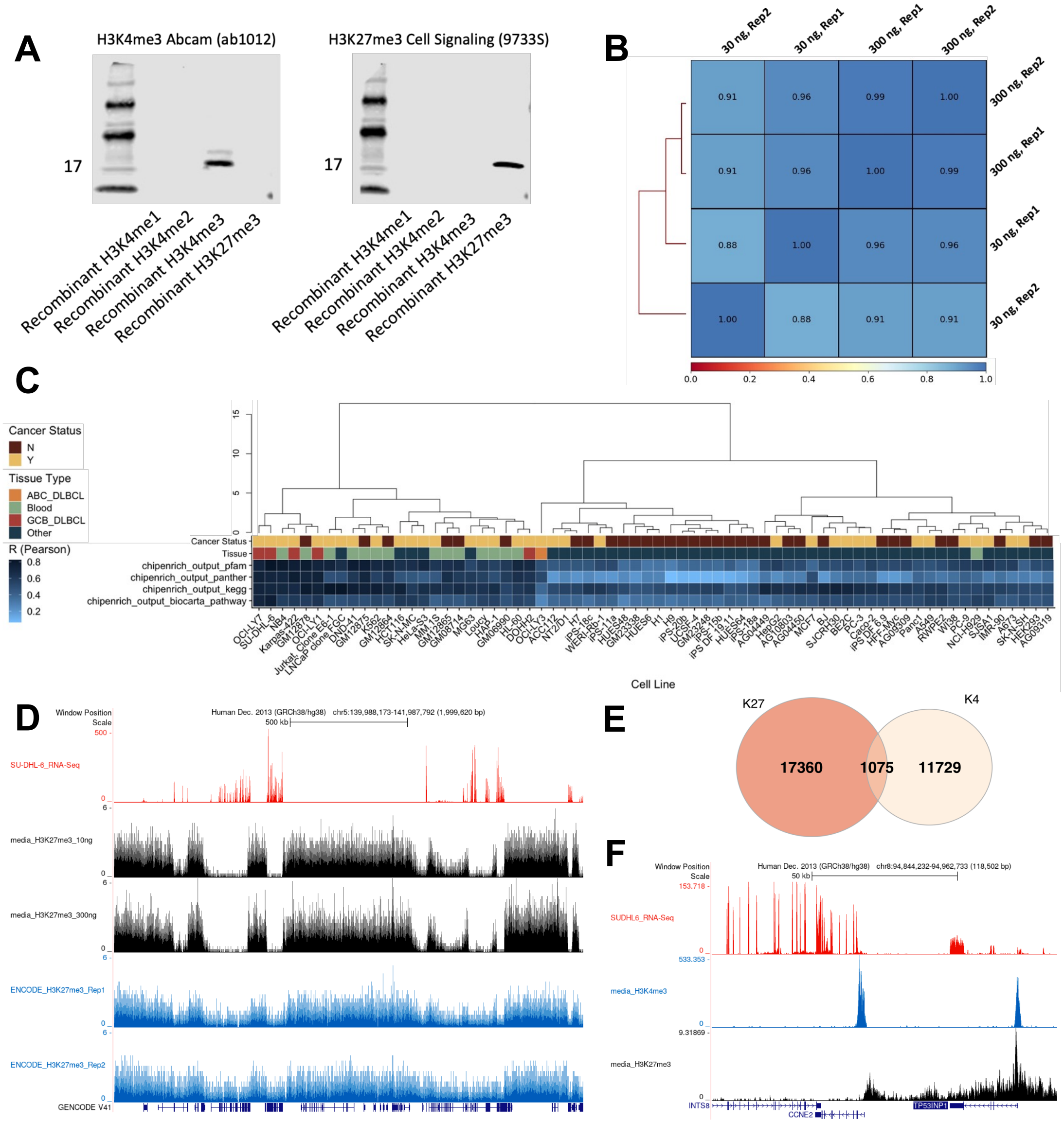

A
H3K4me3 Abcam (ab1012)
H3K27me3 Cell Signaling (9733S)
17
17
Recombinant H3K4me1
Recombinant H3K4me2
Recombinant H3K4me3
Recombinant H3K27me3
Recombinant H3K4me1
Recombinant H3K4me2
Recombinant H3K4me3
Recombinant H3K27me3
B
30 ng, Rep2
30 ng, Rep1
300 ng, Rep1
300 ng, Rep2
300 ng, Rep2
300 ng, Rep1
30 ng, Rep1
30 ng, Rep2
0.91 0.96 0.99 1.00
0.91 0.96 1.00 0.99
0.88 1.00 0.96 0.96
1.00 0.88 0.91 0.91
0.0 0.2 0.4 0.6 0.8 1.0
C
Cancer Status
N
Y
Tissue Type
ABC_DLBCL
Blood
GCB_DLBCL
Other
R (Pearson)
0.8
0.6
0.4
0.2
Cancer Status
Tissue
chipenrich_output_pfam
chipenrich_output_panther
chipenrich_output_kegg
chipenrich_output_biocarta_pathway
Cell Line
D
Window Position
Scale
Human Dec. 2013 (GRCh38/hg38) chr5:139,988,173-141,987,792 (1,999,620 bp)
500 kb
SU-DHL-6_RNA-Seq
media_H3K27me3_10ng
media_H3K27me3_300ng
ENCODE_H3K27me3_Rep1
ENCODE_H3K27me3_Rep2
GENCODE V41
E
K27
K4
17360
1075
11729
F
Window Position
Scale
Human Dec. 2013 (GRCh38/hg38) chr8:94,844,232-94,962,733 (118,502 bp)
50 kb
SUDHL6_RNA-Seq
media_H3K4me3
media_H3K27me3
INTS8
CCNE2
TP53INP1

**Supplementary Figure 6. cfChIP-Seq from SU-DHL-6 conditioned media cf-chromatin reflects distinct chromatin states. A)** Preliminary validation of antibody specificity using recombinant histones (Active Motif) on a western blot. Antibodies against the desired target modifications (H3K4me3 and H3K27me3, respectively) were tested against recombinant H3K4me1, H3K4me2, H3K4me3, and H3K27me3, demonstrating antibody specificity. Additional data for the specificity of the H3K27me3 antibody (Cell Signaling) is described elsewhere[108]. **B)** Pearson correlation across the coverage profiles from Figure 4B. Correlations were summarised over 1000 bp bins, genome wide. **C)** Using ChIPEnrich, pathway analysis using peaks from SU-DHL-6 media profiles was performed and compared to ENCODE H3K4me3 profiles from various cell types (similar to Figure 4E). Each square in the heatmap represents a comparison between SU-DHL-6 media H3K4me3 and H3K4me3 from another cell line. Pearson correlation R values between odds ratios for particular pathway terms, for the different gene sets (including KEGG, Panther, Biocarta, and PFAM), are shown. Multiple gene sets were used to demonstrate correlations independent of the gene set. **D)** Low and high input cfChIP-Seq for H3K27me3 from SU-DHL-6 media cf-chromatin shown, alongside replicates of SU-DHL-6 H3K27me3 ChIP-Seq from ENCODE[69] and SU-DHL-6 RNA-Seq[44]. BigWig files were RPKM normalized before visualization with the UCSC genome browser[48,109]. Tracks were visualized over a large genomic window on chromosome 5. **E)** Overlap of H3K4me3 and H3K27me3 MACS2 peaks from SU-DHL-6 media cf-chromatin (both generated with 300 ng cf-chromatin as input). Overlapping peaks represent bivalent domains. **F)** Visualization of an example bivalent promoter (TP53INP1) using the UCSC genome browser. Tracks represent RNA-Seq from SU-DHL-6, media H3K4me3 (300 ng input), and media H3K27me3 (300 ng input), respectively. All profiles were RPKM normalized before visualization.

### SUPPLEMENTARY TABLES

| Cell Line | Media |
|---|---|
| CAMA-1 | Dulbecco's Modified Eagle Medium |
| HCT116 | Roswell Park Memorial Institute 1640 |
| SUD-HL-6 | Roswell Park Memorial Institute 1640 |
| MCF-7 | Dulbecco's Modified Eagle Medium |
| A549 | Dulbecco's Modified Eagle Medium |

**Supplemetary Table 1:** Culture media used for cell lines in addition to supplementation with 10% FBS and 1% penicillin-streptomycin solution.

| Target | Orientation | Primer Sequence |
|---|---|---|
| short human LINE-1 | Forward | 5'-TCACTCAAAGCCGCTCAACTAC-3' |
| short human LINE-1 | Reverse | 5'-TCTGCCTTCATTTCGTTATGTACC-3' |
| GAPDH | Forward | 5'-GCC AAT CTC AGT CCC TTC CC-3' |
| GAPDH | Reverse | 5'-TAG TAG CCG GGC CCT ACT TT-3' |
| KAT6B | Forward | 5'-GAA GAG GCG GAC CCA GCG GT-3' |
| KAT6B | Reverse | 5'-TTC CTG CCG GTC ATC TCG CTT-3' |
| SLC22A3 | Forward | 5'-GGA GAG GGT GGA CAG ATT GA-3' |
| SLC22A3 | Reverse | 5'-TCA GCC TTG CTG CTA CAG TG-3' |
| QML_93 | Forward | 5'-CAC TGG TTG TCT TTG CAG GA-3' |
| QML_93 | Reverse | 5'-CCT GGG TCA TAT TGG GAC AC-3' |

**Supplementary Table 2:** RNA primer sequences used for qPCR quantification of short human LINE-1 and ATAC-Seq quality control qPCR for enrichment of open regions (GAPDH and KAT6B) relative to closed regions (SLC22A3 and QML_93).

| Source | ID | Cell Type/Tissue (Epigenome) |
|---|---|---|
| REMC | E003 | H1_Cell Line |
| REMC | E004 | H1_BMP4_Derived_Mesendoderm_Cultured_Cells |
| REMC | E005 | H1_BMP4_Derived_Trophoblast_Cultured_Cells |
| REMC | E006 | H1_Derived_Mesenchymal_Stem_Cells |
| REMC | E007 | H1_Derived_Neuronal_Progenitor-Cultured_Cells |
| REMC | E011 | hESC_Derived_CD184+_Endoderm_Cultured_Cells |
| REMC | E012 | hESC_Derived_CD56+_Ectoderm_Cultured_Cells |
| REMC | E013 | hESC_Derived_CD56-_Mesoderm_Cultured_Cells |
| REMC | E016 | HUES64_Cell Line |
| REMC | E024 | 4star |
| REMC | E027 | Breast_Myoepithelial_Cells |
| REMC | E028 | Breast_vHMEC |
| REMC | E037 | CD4_Memory_Primary_Cells |
| REMC | E038 | CD4_Naive_Primary_Cells |
| REMC | E047 | CD8_Naive_Primary-Cells |
| REMC | E050 | Mobilized_CD34_Primary_Cells_Female |
| REMC | E053 | Neurosphere_Cultured_Cells_Cortex_Derived |
| REMC | B054 | Neurosphere_Cultured_Cells_Ganglionic_Eminence_Derived |
| REMC | E055 | Penis_Foreskin_Fibroblast_Primary_Cells_skin01 |
| REMC | B056 | Penis_Foreskin_Fibroblast_Primary_Cells_skin02 |
| REMC | E057 | Penis_Foreskin_Keratinocyte_Primary_Cells_skin02 |
| REMC | E058 | Penis_Foreskin_Keratinocyte_Primary_Cells_skin03 |
| REMC | E059 | Penis_Foreskin_Melanocyte_Primary_Cells_skin01 |
| REMC | BO61 | Penis_Foreskin_Melanocyte_Primary_Cells_skin03 |
| REMC | E062 | Peripheral_Blood_Mononuclear_Primary_Cells |
| REMC | E065 | Aorta |
| REMC | E066 | Adult Liver |
| REMC | E070 | Brain_Germinal_Matrix |
| REMC | E071 | Brain_Hippocampus_Middle |
| REMC | E079 | Esophagus |
| REMC | E082 | Fetal_Brain_Female |
| REMC | BO84 | Fetal_Intestine_Large |
| REMC | E085 | Fetal Intestine_Small |
| REMC | E087 | Pancreatic_Islets |
| REMC | E094 | Gastric |
| REMC | E095 | Left_Ventricle |
| REMC | E096 | Lung |
| REMC | E097 | Ovary |

| | | |
|---|---|---|
| REMC | E098 | Pancreas |
| REMC | E100 | Psoas_Muscle |
| REMC | E104 | Right_Atrium |
| REMC | E105 | Right_Ventricle |
| REMC | E106 | Sigmoid_Colon |
| REMC | E109 | Small_Intestine |
| REMC | E112 | Thymus |
| REMC | E113 | Spleen |
| REMC | E114 | A549 |
| REMC | E116 | GM12878 |
| REMC | E117 | HELA |
| REMC | D118 | HEPG2 |
| REMC | E119 | HMEC |
| REMC | E120 | HSMM |
| REMC | E122 | HUVEC |
| REMC | E123 | K562 |
| REMC | E127 | NHEK |
| REMC | E128 | NHLE |

**Supplementary Table 3:** 56 cell types and tissues used in this study from the REMC database.

| Source | H3K4me3 Experiment ID | H3K27me3 Experiment ID(s) | Cell Line |
|---|---|---|---|
| ENCODE | ENCFF361TGO | | A549 |
| ENCODE | ENCFF272QER | | A673 |
| ENCODE | ENCFF081HTX | | ACC112 |
| ENCODE | ENCFF824TEY | | AG04449 |
| ENCODE | ENCFF988TOJ | | AG04450 |
| ENCODE | ENCFF949SYV | | AG09309 |
| ENCODE | ENCFF643VVG | | AG09319 |
| ENCODE | ENCFF313CEA | | AG10803 |
| ENCODE | ENCFF251CDW | | BE2C |
| ENCODE | ENCFF364DKX | | BJ |
| ENCODE | ENCFF642BGI<br>ENCFF001WWT,ENCFF876QHF | ENCFF001WWL,ENCFF604KXO | Caco-2 |
| ENCODE | ENCFF069RHJ | | DND-41 |
| ENCODE | ENCFF375YDP | | DOHH2 |
| ENCODE | ENCFF253XQQ | | GM06990 |
| ENCODE | ENCFF800MHH | | GM08714 |
| ENCODE | ENCFF704LTU | | GM12864 |
| ENCODE | ENCFF438JMP | | GM12865 |
| ENCODE | ENCFF205BPF | | GM12875 |
| ENCODE | ENCFF320OGZ<br>ENCFF001WYJ,ENCFF795URC | ENCFF001WYB,ENCFF247VUO | GM12878 |
| ENCODE | ENCFF123ETI | | GM23248 |
| ENCODE | ENCFF387WKX | | GM23338 |
| ENCODE | ENCFF041HYH<br>ENCFF001SVC,ENCFF192QQV | ENCFF001SUY,ENCFF434CYZ | H1 |
| ENCODE | ENCFF985GWM<br>ENCFF001XAR,ENCFF207YLR | ENCFF001WZT,ENCFF302BSC | H7 |
| ENCODE | ENCFF473AUA | | H9 |
| ENCODE | ENCFF856YLE | | HAP-1 |
| ENCODE | ENCFF187LLD | | HCT116 |
| ENCODE | ENCFF617NUV | | HEK293 |

| | | | |
|---|---|---|---|
| ENCODE | ENCFF469OXD | | HeLa-S3 |
| ENCODE | ENCFF549DKP<br>ENCFF001XDB,ENCFF712HMU | ENCFF001XCT,ENCFF042EDV | HepG2 |
| ENCODE | ENCFF366OCN | | HFF-Myc |
| ENCODE | ENCFF021JBH | | HL-60 |
| ENCODE | ENCFF441QBU | | HUES48 |
| ENCODE | ENCFF023OID | | HUES6 |
| ENCODE | ENCFF187LIH | | HUES64 |
| ENCODE | ENCFF093NQC | | IMR-90 |
| ENCODE | ENCFF450VVJ | | iPS DF 19.11 |
| ENCODE | ENCFF150NDM | | iPS DF 6.9 |
| ENCODE | ENCFF454SUG | | iPS-11a |
| ENCODE | ENCFF724ZXV | | iPS-15b |
| ENCODE | ENCFF618IAG | | iPS-18a |
| ENCODE | ENCFF699VZO | | iPS-18c |
| ENCODE | ENCFF483YLD | | iPS-20b |
| ENCODE | ENCFF221IPH | | Jurkat, Clone E6-1 |
| ENCODE | ENCFF122CSI<br>ENCFF001XGT,ENCFF378OQB | ENCFF001SZF,ENCFF126QYP | K562 |
| ENCODE | ENCFF693UVS<br>[ENCFF125FKI] | [ENCFF840XGT] | Karpas-422 |
| ENCODE | ENCFF586HAM | | LNCaP clone FGC |
| ENCODE | ENCFF329ZUR | | Loucy |
| ENCODE | ENCFF145CCI | | MCF7 |
| ENCODE | ENCFF586UCO | | MG63 |
| ENCODE | ENCFF367SUW | | MM.1S |
| ENCODE | ENCFF527YWO | | NB4 |
| ENCODE | ENCFF577HWW | | NCI-H929 |
| ENCODE | ENCFF954FXD | | NT2/D1 |
| ENCODE | ENCFF670JNI | | OCI-LY1 |
| ENCODE | ENCFF826PSG<br>[ENCFF224FBU] | [ENCFF254CXL,ENCFF375IRO] | OCI-LY3 |

| | | | |
|---|---|---|---|
| ENCODE | ENCFF392YEW | | OCI-LY7 |
| ENCODE | ENCFF103AWU | | Panc1 |
| ENCODE | ENCFF375RGR | | PC-3 |
| ENCODE | ENCFF347EMU | | PC-9 |
| ENCODE | ENCFF965NTW | | RWPE2 |
| ENCODE | ENCFF639GVZ | | SJCRH30 |
| ENCODE | ENCFF870NJQ | | SJSA1 |
| ENCODE | ENCFF337FTH | | SK-N-MC |
| ENCODE | ENCFF682JYE | | SK-N-SH |
| ENCODE | ENCFF884HJC [ENCFF408JOT] | [ENCFF182RDK] | SU-DHL-6 |
| ENCODE | ENCFF742FZZ | | UCSF-4 |
| ENCODE | ENCFF287NSO | | WERI-Rb-1 |
| ENCODE | ENCFF033PCY | | WI38 |

**Supplementary Table 4:** H3K4me3 and H3K27me3 profiles from 68 cell lines used in this study, from the ENCODE database. If comma separated, samples indicate broad and narrow peaks, respectively. If accession ID is enclosed by square brackets, further processing of replicate BAM files was performed to broad and narrow peak file format.
